# Choreography of human mitochondrial leaderless mRNA translation initiation

**DOI:** 10.1101/2025.07.10.662049

**Authors:** Shuangjie Shen, Yinghua Xu, Daniel L. Kober, Jinfan Wang

## Abstract

It remains unclear how human mitochondrial ribosomal subunits assemble into an elongation-competent 55S particle on mRNAs devoid of 5’ leader sequences. Here, we reconstituted and directly tracked human mitochondrial translation initiation using real-time single-molecule fluorescence spectroscopy. Corroborated with cryo-EM structural analysis, we show that the initiation factor mtIF2 and initiator fMet-tRNA^Met^ are loaded to the 28S subunit to drive mRNAs binding via 5’ start codon recognition. This enables sequential loading of the two ribosomal subunits onto the leaderless mRNA to initiate. In parallel, a preassembled 55S monosome can also be loaded with mtIF2 and fMet-tRNA^Met^ to initiate on the mRNA. Both initiation pathways yield active complexes to enter translation elongation, which is gated by mtIF2. The monosome loading pathway can initiate promiscuously with non-formylated Met-tRNA^Met^, thus its usage may under tight regulation in cells, e.g. by mtIF3. Our work provides a dynamic framework for the distinct human mitochondrial translation initiation.

## Introduction

To initiate protein synthesis, the ribosomes are recruited to bind the mRNA, forming an active initiation complex with the initiator tRNA recognizing a start codon programmed in the ribosomal P site. This process is tightly regulated to control the identity and quantity of the synthesized proteins for cellular homeostasis^1^. Despite decades of studies^2^, the dynamic mechanism of mammalian (including human) mitochondrial translation initiation is poorly understood. The mammalian mitochondrial ribosomes are considerably divergent from the bacterial or cytosolic ribosomes. Their ribosomal RNA content is reduced by ∼1 MDa in molecular mass, which is partially compensated by acquiring mitochondria-specific ribosomal proteins or else by incorporating extension and insertions into otherwise conserved ribosomal proteins^3,4^. The mRNA binding channel on the 28S small subunit is also fundamentally remodeled (Figure S1), with the uS5m protein forming a latch that connects the head and the body of the 28S subunit^3,4^, presenting an extra physical barrier at the mRNA entry path for mRNA loading. How the mitochondrial mRNAs are loaded onto the ribosomes to initiate protein synthesis remains an open question^5^. The mitochondrial mRNAs do not possess a 7-methylguanosine cap at the 5’ end as in the cytoplasmic mRNAs to stimulate ribosome binding and scanning for a start site^1,6,7^. Neither do they contain the classical bacterial Shine-Dalgarno sequence in 5’ untranslated regions (UTRs) to guide high-affinity binding to the small ribosomal subunit with the start codon directly placed in the P site^8–11^. In fact, mammalian mitochondrial mRNAs lack a 5’ UTR, or else have only a very short one^12^, and thus are referred to as leaderless mRNAs.

It has been established that leaderless mRNA translation initiation is facilitated by *N*-formylmethionine-tRNA^Met^ (fMet-tRNA^Met^) and two protein factors (mtIF2 and mtIF3)^2,13–15^. Recent studies observed binding of leaderless mRNAs and fMet-tRNA^Met^ to the human mitochondrial 55S monosomes, but not to the 28S small ribosomal subunit^16,17^. A monosome-loading mechanism was thus proposed as a distinct feature for mitochondrial leaderless mRNA translation initiation^16,17^. By contrast, earlier studies showed that leaderless mRNAs can bind mammalian 28S subunits *in vitro*^18,19^ and *in vivo*^20^, implicating the possibility of sequentially loading the 28S small and 39S large ribosomal subunits onto a leaderless mRNA for initiation^18,21^. These discrepancies may arise from the limitations of traditional methods to deconvolute heterogenous pathways with temporal resolution^2^. We thus combined real-time single-molecule spectroscopy and single-particle cryo-EM approaches to unravel the dynamic molecular choreography of mitochondrial translation initiation.

## Results

### Imaging of leaderless mRNA binding to mitochondrial ribosomes

Here, we reconstituted human mitochondrial translation initiation *in vitro*^16,22–25^ (Figure S2) and established a real-time single-molecule assay to directly monitor leaderless mRNA binding to human mitochondrial ribosomes (Figure 1A). The 28S small subunit was site-specifically biotinylated via an A1-tag^26^ fused to the C-terminus of mS26 (biotin-28S, Figures S2A-S2D)^26^. The A1-tagged mS26 fully supported mitoribosome activities in cells as demonstrated by an on-gel mito-FUNCAT assay (Figure S3)^27^. Before testing full-length natural leaderless mRNAs, we examined a 45-nt model mRNA that contained a single AUG codon at the monophosphorylated 5’ end and a Cy3 dye at the 3’ end, avoiding ambiguities in start site usage.

**Figure 1.**
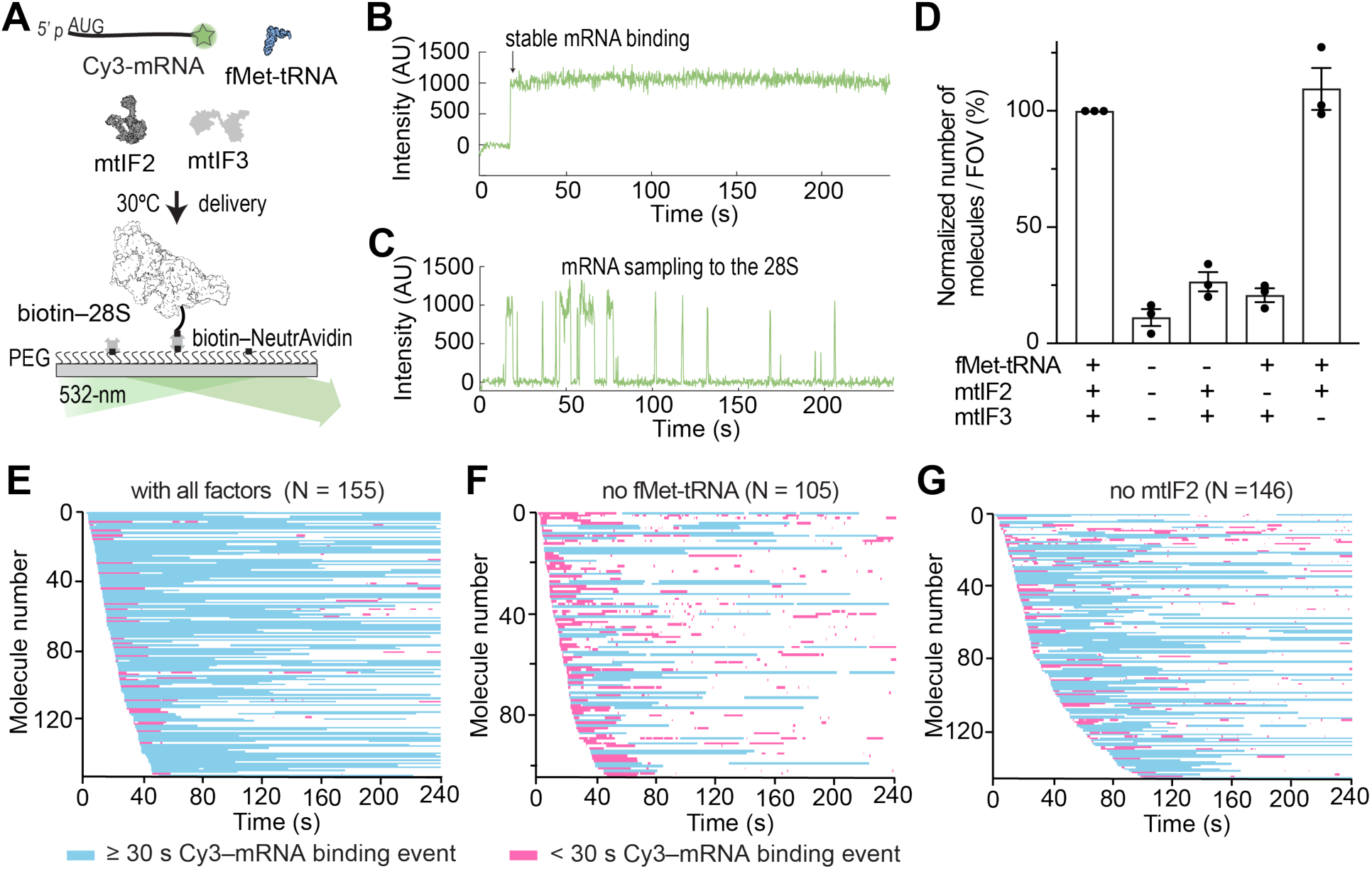
Single-molecule assay for observing mRNA–28S subunit binding. **(A)** Schematic of assay to track Cy3-mRNA binding to tethered 28S subunits. **(B)** Example fluorescence trace of showing stable binding of mRNA (Cy3, green) to the 28S subunit. **(C)** Example fluorescence trace of mRNA sampling to the 28S subunit. **(D)** Mean normalized number of Cy3 molecules observed per field of view (FOV) in reactions under varying conditions. Indicated errors are standard error of the mean from three biological replicates with points representing the value from individual replicates. **(E-G)** Stacks of single-molecule traces from different reaction conditions, sorted by the arrival of the first Cy3 binding event. Each row represents a single 28S complex. Cy3-mRNA binding events are display in blue (≥ 30 s) or pink (< 30 s). N indicates number of analyzed molecules.

We first asked whether the model mRNA can be loaded onto the 28S subunit in the absence of the 39S subunit. Accordingly, biotin-28S was tethered to neutravidin-coated glass imaging surface (Figure 1A). Upon start of real-time data acquisition on a total internal reflection fluorescence microscope (TIRFM) with a 532-nm laser excitation, a mixture of 1 nM Cy3 model mRNA, 200 nM fMet-tRNA^Met^, 400 nM mtIF2, 300 nM mtIF3, and excess GTP were delivered to the flow chamber at 30°C by a stopped-flow pump. Notably, binding of the model leaderless mRNA to the 28S subunit was readily detected as the appearance of Cy3 signals upon sample mixing (Figures 1B and 1C), and we normalized the number of observed Cy3 molecules per field of view (FOV) as 100% mRNA binding efficiency (Figures 1D and S4A). The majority of the mRNA binding events were stable (lifetime > 30 s) (Figures 1B and 1E), while some showed transient sampling of the mRNA to the 28S subunit (Figures 1C, 1E and S4B). The mRNA binding efficiency was reduced by 97% when the surface was pretreated with biotin to block biotin-28S immobilization (Figure S4A), assuring that the Cy3 binding events were specific to the tethered 28S subunits. These results indicated that the model leaderless mRNA can indeed bind the 28S subunits.

We next repeated the experiments with full-length, naturally occurring leaderless Cox3 and CytB mRNAs that were labeled with Cy3 at the 3’ end. Here, the mRNA binding efficiencies were ∼77% and ∼44% for Cox3 and CytB mRNAs, respectively, and most of the mRNA binding events were stable (Figures S4A, S4B, S4G and S4H). These results suggested that the stable assembly of a leaderless mRNA–28S complex is a generalizable feature of human mitochondrial translation initiation, consistent with earlier biochemical findings^18,19^.

### Stable mRNA binding requires mtIF2 and start codon recognition by fMet-tRNA^Met^

We next sought to determine the factor requirement for leaderless mRNA binding to the 28S subunit. When only the model leaderless mRNA was delivered to the 28S subunits, the mRNA binding efficiency was ∼11% and the resulting binding events were largely in the transient sampling states (Figures 1D and S4A-S4C). Similar results were observed when either fMet-tRNA^Met^ or mtIF2 was omitted from the reaction (mRNA binding efficiency ∼26.5% or 20.8%, respectively, Figures 1D, 1F, 1G, S4A, and S4B). On the other hand, the absence mtIF3 did not inhibit mRNA binding nor disrupt the mRNA–28S complex stability (Figures 1D, S4A, S4B, and S4D). Thus, mRNA binding is driven by mtIF2 and fMet-tRNA^Met^.

To test whether a start codon at the 5’ end determines mRNA binding, we mutated the AUG to a non-cognate CAA codon (no-AUG mRNA). With this no-AUG mRNA, the mRNA binding efficiency was reduced to ∼11.6% (Figures S4A, S4B, and S4E). Remarkably, when a 29-nucleotides 5’ UTR was added upstream of the AUG codon (29nts-UTR mRNA), the mRNA binding efficiency was limited to ∼9.3% (Figures S4A, S4B, and S4F). Thus, efficient and stable mRNA binding to the 28S subunits requires a start codon at the 5’ end being recognized by the fMet-tRNA^Met^ in the presence of mtIF2.

### The 28S subunit is loaded with mtIF2 and fMet-tRNA^Met^ to recruit mRNAs

We next sought to determine the relative timing of mRNA and factor binding to the 28S subunit. In the presence of all factors, the association of the Cy3 model mRNA with the 28S subunit went to completion by ∼1 min (Figure 1E), with the observed association kinetics fits better to a two-step reaction model than a single-step model (Figure S4I). These results suggested that the assembly of a mRNA–28S complex was a multi-step process. We hypothesized that the first rate-limiting step could be pre-loading the 28S subunit with mtIF2 and fMet-tRNA^Met^ that are critical for mRNA binding (Figures 1F and 1G).

We thus fluorescently labeled mtIF2 with Cy5 at the flexible N-terminus to track the dynamics of this factor. We also moved the Cy3 label on the mRNA to the 14^th^ nucleotide position from the 5’ end (mRNA-Cy3), which was predicted to be within FRET distance (∼50 Å) to the mtIF2 N-terminus when both were bound on the 55S monosome (Figure S5A)^22^. Alternative laser excitation (ALEX)^28^ at wavelengths of 532 and 640 nm was employed here: the 640-nm laser can directly excite Cy5-mtIF2 to enable direct monitoring of mtIF2, while the 532-nm laser excitation enables FRET analysis between mtIF2 and the mRNA to score for functional complex assemblies (Figure 2A). At the beginning to data acquisition, mRNA-Cy3 and Cy5-mtIF2 were co-delivered with fMet-tRNA^Met^ and mtIF3 to immobilized 28S subunits. Analysis of directly excited Cy5 (mtIF2) and Cy3 (mRNA) fluorescence intensities (Figure 2B), aligned to the beginning of the Cy5 events, demonstrated that Cy5-mtIF2 binding occurred prior to that of the mRNA (Figure 2C). Immediately when the mRNA was loaded, a stable mRNA–mtIF2 FRET event was detected (Figure 2B), which was further validated in experiments using a single 532-nm laser excitation (Figures S5B-S5D). Both mtIF2 binding and the subsequent mRNA loading to the 28S were concentration-dependent, and the association kinetics were fit to a single-exponential function with pseudo-second order rate constants of *∼*4.8 and *∼*21.6 µM^-1^s^-1^, respectively (Figures 2D-2G). Omission of mtIF3 had a minor impact on the dynamics of mtIF2 and mRNA binding (Figures 2D-2G), consistent with mtIF3 being dispensable for mRNA–28S subunit binding observed above (Figures 1D and S4D). When non-formylated Met-tRNA^Met^ was used in place of the fMet-tRNA^Met^, a different mRNA–mtIF2 FRET behavior was observed, where mtIF2 transiently cycled on and off the complex (Figure S5E-S5G). Thus, the *N*-formylation of Met-tRNA^Met^ plays an important role in its recognition by mtIF2, which in turn is required for stable mtIF2 binding on the 28S subunit.

**Figure 2.**
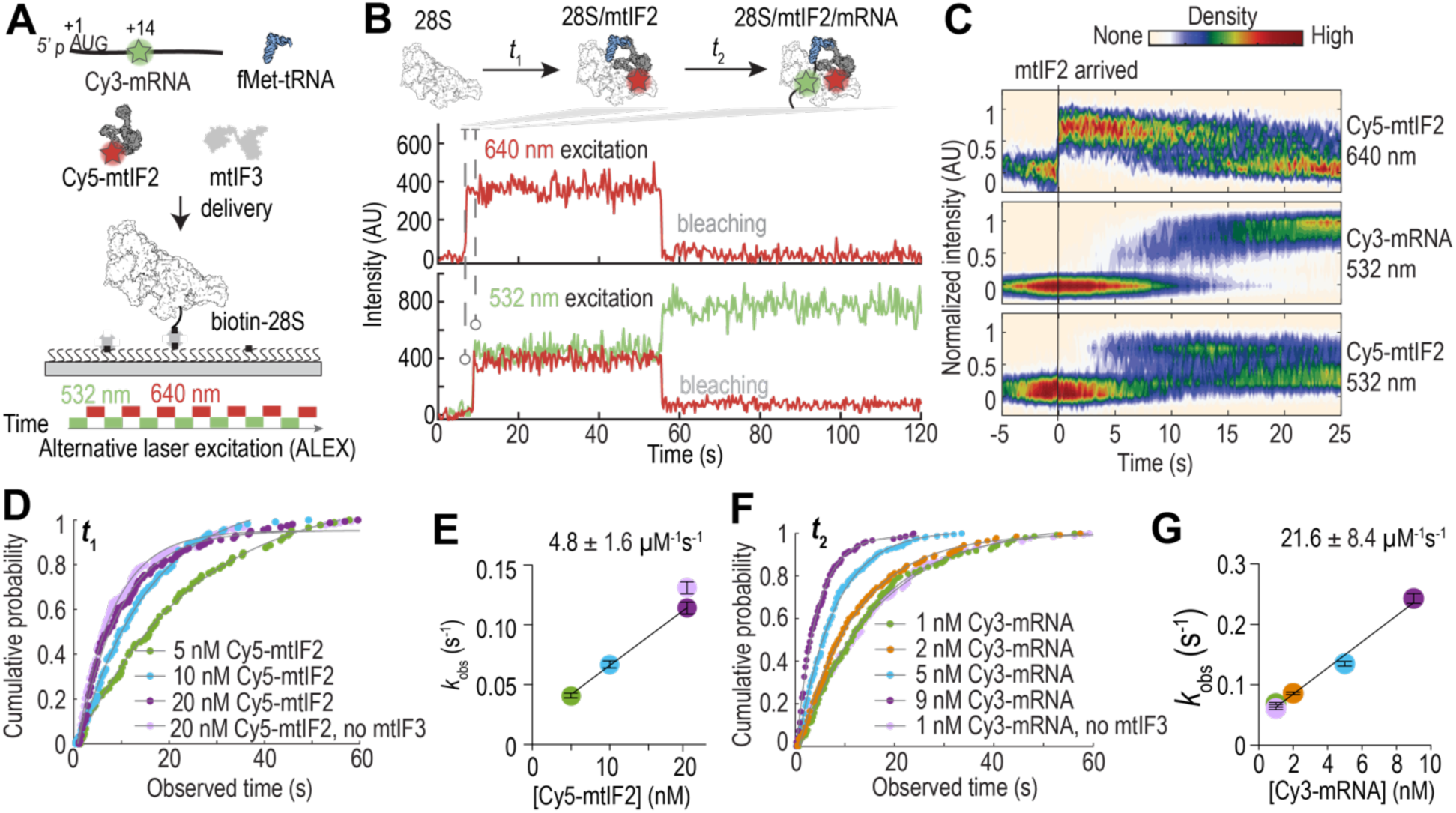
Observing mtIF2 and mRNA binding to 28S subunits. **(A)** Schematic of assay to track Cy3-mRNA and Cy5-mtIF2 binding to tethered 28S subunits using alternative laser excitation. **(B)** Example fluorescence trace of mtIF2 (Cy5, red) and mRNA (Cy3, green) binding to the 28S subunit. Signals from the 640-nm (middle) or 532-nm (bottom) laser excitation movie frames are shown. **(C)** Density heat map of the normalized fluorescence intensities of Cy5-mtIF2 and Cy3-mRNA from direct excitation and Cy5-mtIF2 from FRET synchronized to the binding of mtIF2 to the 28S subunit (N = 158). **(D)** Binding of mtIF2 to 28S subunits is concentration-dependent; arrival time distributions were fit to a single-exponential model. **(E)** Linear regression of observed rates of mtIF2 binding (*k*_obs_) estimated from (**D**). **(F)** Binding of mRNA to mtIF2-bound 28S complex is concentration-dependent; arrival time distributions were fit to a single-exponential model. **(G)** Linear regression of observed rates of mRNA binding (*k*_obs_) estimated from (**F**). Indicated errors are 95% confidence intervals.

The requirement of fMet recognition for stable mtIF2 binding prompted us to determine the relative timing of fMet-tRNA^Met^ arrival. To enable direct tracking of the tRNA, we labeled the fMet-tRNA^Met^ at the flexible U17 with Cy5 via a chemically-installed C6-NH_2_ linker (Figure S6A). Such synthetic tRNAs have been used in studies of yeast and human cytosolic translation^29,30^. After tethering biotin-28S on the imaging surface, a mixture of Cy5-fMet-tRNA^Met^, Cy5.5-mtIF2, mRNA-Cy3, and mtIF3 was delivered with data acquisition performed under ALEX excitation with the 532 and 640 nm lasers (Figure 3A). Binding of Cy5.5-mtIF2 and Cy5-fMet-tRNA^Met^ was monitored by direct 640-nm laser excitation, and functional complex assembly was demonstrated by the FRET between mRNA-Cy3 and Cy5.5-mtIF2. For most of the molecules (∼94%, 99 out of 105), we observed sequential association of mtIF2, fMet-tRNA^Met^ and the mRNA with the 28S subunit (Figure 3B). For the rest of the molecules (∼6%, 6 out of 105), mtIF2 and fMet-tRNA^Met^ co-arrived before the mRNA binding (Figure S6B). At a constant Cy5.5-mtIF2 concentration (10 nM), the Cy5-fMet-tRNA^Met^ and mRNA–Cy3 association kinetics were concentration-dependent (∼5 and ∼31 µM^-1^s^-1^, respectively; Figures 3D-3G). The fraction of molecules that showed simultaneous mtIF2 and fMet-tRNA^Met^ binding increased (∼16%, 16 out of 101) when the Cy5-fMet-tRNA^Met^ concentration was doubled. These co-arrival events likely reflected mtIF2 and fMet-tRNA^Met^ preforming a complex before associating with the 28S subunit^22,31^. Once loaded, fMet-tRNA^Met^ remained stably bound on the complex (median lifetime ∼67 s, Figure S6C), suggesting the formation of a stable 28S–mtIF2–fMet-tRNA^Met^–mRNA pre-initiation complex. Thus, mtIF2 and fMet-tRNA^Met^ are recruited to bind the 28S subunit to stimulate rapid and stable loading of leaderless mRNAs.

**Figure 3.**
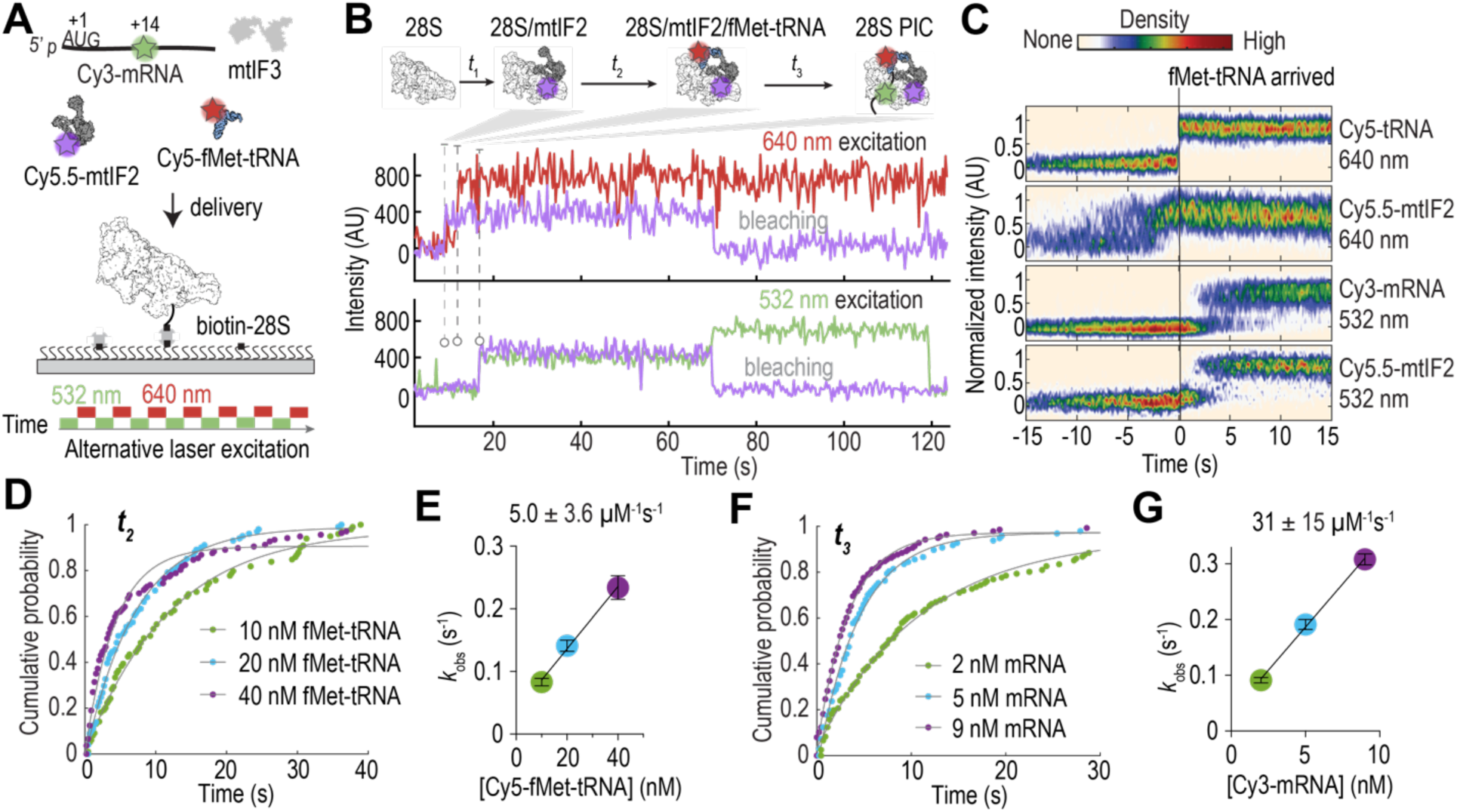
Direct tracking of fMet-tRNA^Met^. **(A)** Schematic of assay to track Cy3-mRNA, Cy5.5-mtIF2, and Cy5-fMet-tRNA^Met^ binding to tethered 28S subunits using alternative laser excitation. **(B)** Example fluorescence trace of mtIF2 (Cy5.5, purple), fMet-tRNA^Met^ (Cy5, red) and mRNA (Cy3, green) binding to the 28S subunit. Signals from the 640-nm (middle) or 532-nm (bottom) laser excitation movie frames are shown. **(C)** Density heat map of the normalized fluorescence intensities of Cy5-fMet-tRNA^Met^, Cy5.5-mtIF2 and Cy3-mRNA from direct excitation and Cy5.5-mtIF2 from FRET synchronized to the binding of fMet-tRNA^Met^ to the 28S subunit (N = 105). **(D)** Binding of fMet-tRNA^Met^ to mtIF2-bound 28S complexes is concentration-dependent; arrival time distributions were fit to a single-exponential model. **(E)** Linear regression of observed rates of fMet-tRNA^Met^ binding (*k*_obs_) estimated from (**D**). **(F)** Binding of mRNA to 28S–mtIF2–fMet-tRNA^Met^ complex is concentration-dependent; arrival time distributions were fit to a single-exponential model. **(G)** Linear regression of observed rates of mRNA binding (*k*_obs_) estimated from (**F**). Indicated errors are 95% confidence intervals.

### Structure of the 28S–mtIF2–fMet-tRNA^Met^–mRNA complex

To provide structural evidence of leaderless mRNAs binding on the 28S subunit and to understand how mtIF2 and fMet-tRNA^Met^ collaborate to promote mRNA binding, we designed a vitrified sample to solve the structure of the 28S–mtIF2–fMet-tRNA^Met^–mRNA complex (28S PIC) by single-particle cryo-EM analysis (Figure S7). Model leaderless mRNAs were added to a mixture of 28S subunits, mtIF2•GTP, mtIF3, and fMet-tRNA^Met^ at room temperature before application to a cryo-EM grid and plunge freezing. The time elapsed from sample mixing to vitrification was ∼1 min. At this pre-steady state timepoint, we expect that the mRNA binding should have occurred on a majority of the 28S subunits (Figure S4J), resulting in a stable complex with mtIF2 (Figure S5D) and fMet-tRNA^Met^ (Figure S6C). Indeed, after data processing, we identified a subset of around 167,631 particles (representing ∼57% of all 28S particles) containing well defined densities for mtIF2 and fMet-tRNA^Met^ (Figure S7D). These particles belonged to three classes in three-dimensional classification, with two of these classes combined as the final set of 107,666 particles that were refined to a 2.50 Å map (28S PIC) (Figures S7E and S7F).

In the 28S PIC map, the fMet-tRNA^Met^ and mtIF2 were better resolved in the regions that contacted the 28S subunit, including the anticodon arm of the tRNA^Met^ and domains II and III of mtIF2 (Figures 4A and S7G). The solvent-exposed acceptor stem of tRNA^Met^ and its interacting domain IV of mtIF2 were less well resolved, suggesting conformational flexibilities in these peripheral regions of the complex (Figure S7G). Densities that belong to the mRNA can be traced from the P site to the mS39 region in the 3’ direction (Figure 4B), but only the first three nucleotides that corresponded to the AUG codon gave densities strong enough for confident modeling (Figures 4C and 4D). A close examination near the P site region showed that the mRNA densities started with the 5’ monophosphate group, followed by the AUG codon that formed strong base-pairing interactions with the anticodon of the fMet-tRNA^Met^ (Figure 4D). These results suggested that the codon–anticodon interactions played a major role in stabilizing the mRNA on the ribosome, and thus provided the molecular basis for the strict dependence of leaderless mRNA binding on the 5’ start codon recognition by fMet-tRNA^Met^. The 28S PIC structure also revealed how mtIF2 stimulates mRNA binding via its direct contacts with the fMet-tRNA^Met^ and the 28S subunit.

**Figure 4.**
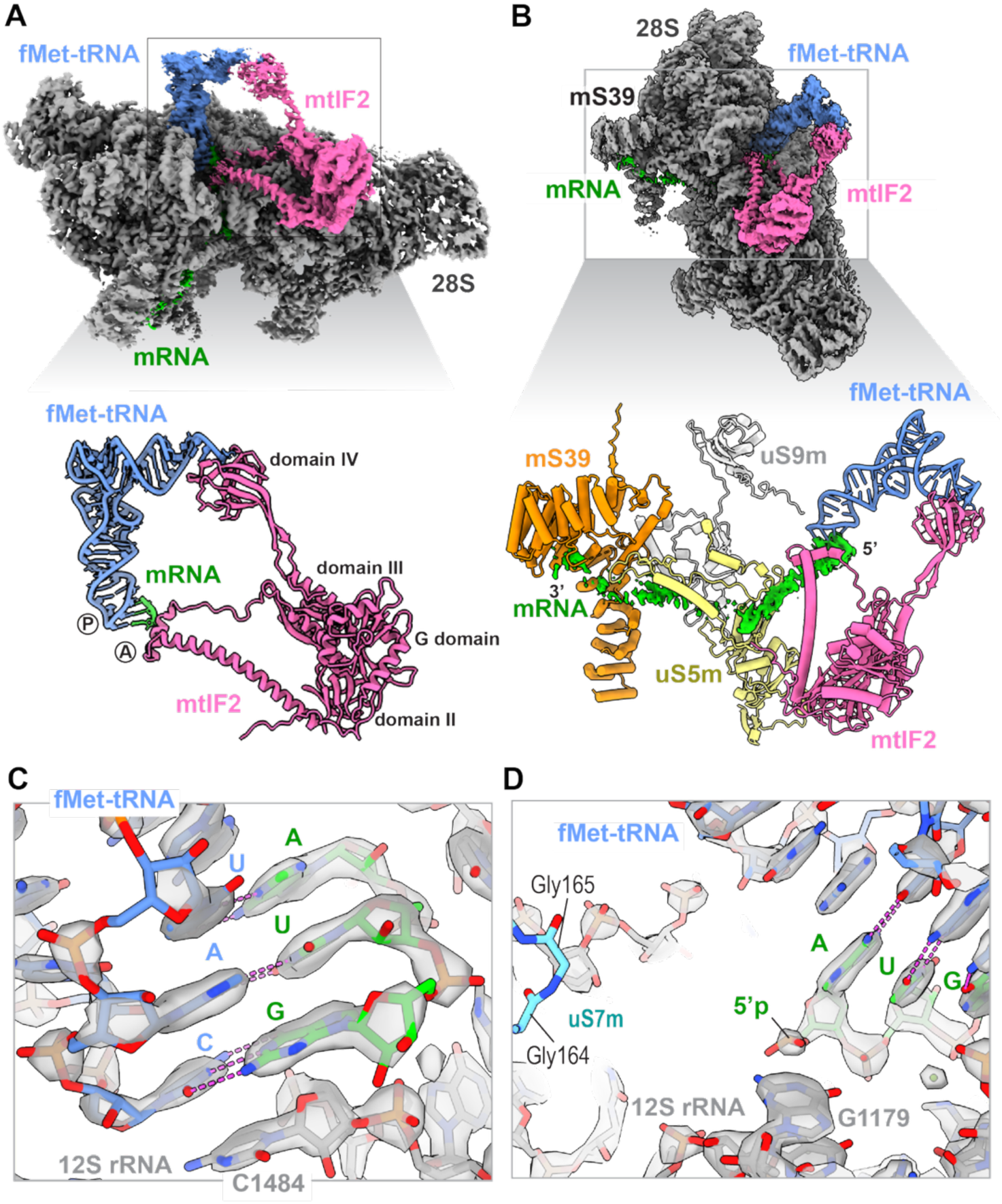
Cryo-EM structure of the 28S PIC. **(A)** Cryo-EM density map of the 28S PIC (top, overall resolution 2.5 Å) and the domain structure of mtIF2 (bottom) in the context of fMet-tRNA^Met^ and mRNA interactions. In the map, the 28S subunit is colored in gray, fMet-tRNA^Met^ in blue, mtIF2 in pink and mRNA in green. The A and P site regions are indicated in the model. **(B)** A side view of the 28S PIC cryo-EM map (top) and a detailed view of the structural model with the denoted components displayed in isolation (bottom) showing the mRNA path. **(C)** A detailed view showing the codon–anticodon interactions with the cryo-EM density shown in transparent surface. **(D)** A detailed view surrounding the 5’ end of the mRNA with the cryo-EM density shown in transparent surface.

### Two parallel pathways of ribosome loading onto leaderless mRNAs

Having established the assembly of a 28S PIC in the absence of the 39S subunit, we hypothesized that the two ribosomal subunits should be able to load onto a leaderless mRNA sequentially. To test this, we designed an inter-ribosomal subunit FRET signal to track 55S monosome assembly (Figure S2G). The S6 peptide tag^26^ was appended to endogenous mS31 (28S subunit) or bL31m (39S subunit) proteins at the C-termini, which are within predicted FRET distance (∼40Å) in structural models of 55S ribosomes (Figures S2A and S2G-S2I). The tagged ribosomes were functional in cells (Figure S3), and purified 28S-S6 and 39S-S6 subunits were site-specifically labeled with Cy3 and Cy5 fluorescent dyes, respectively (Figure S2J).

To observe ribosome loading, the model leaderless mRNA was biotinylated at the 3’end and tethered to the imaging surface (Figure 5A). Upon start of data acquisition, a mixture with 1.5 Cy3-28S, 4 nM Cy5-39S, 400 nM mtIF2, 200 nM fMet-tRNA^Met^ and 300 nM mtIF3 was added to the flow chamber. When excited by a 532-nm laser, the appearance of Cy3 fluorescence indicated Cy3-28S binding to the mRNA, while the appearance of Cy3–Cy5 FRET indicated the formation of a 55S initiation complex (IC) (Figures 5B, 5C, and S8). Under this condition, the majority of the observed 55S IC (∼88%) were assembled by loading the 28S subunit first, followed by the joining of the 39S subunits (Figures 5B, 5D, and 5E). These results were consistent with the formation of a 28S PIC on the leaderless mRNA and confirmed that the 28S PIC was active in full initiation. Another small fraction of the molecules (∼12%) showed simultaneous binding of the two ribosomal subunits to the mRNA in a FRET state, indicating direct monosome loading (Figures 5C, 5D, and 5E). Similar results were obtained with full-length Cox3 and CytB mRNAs, although with varying pathway frequencies and kinetics (Figures 5D and 5E). These results demonstrated that leaderless mRNA translation initiation can take heterogeneous pathways, with the mitochondrial ribosomal subunits loading sequentially or simultaneously as a monosome.

**Figure 5.**
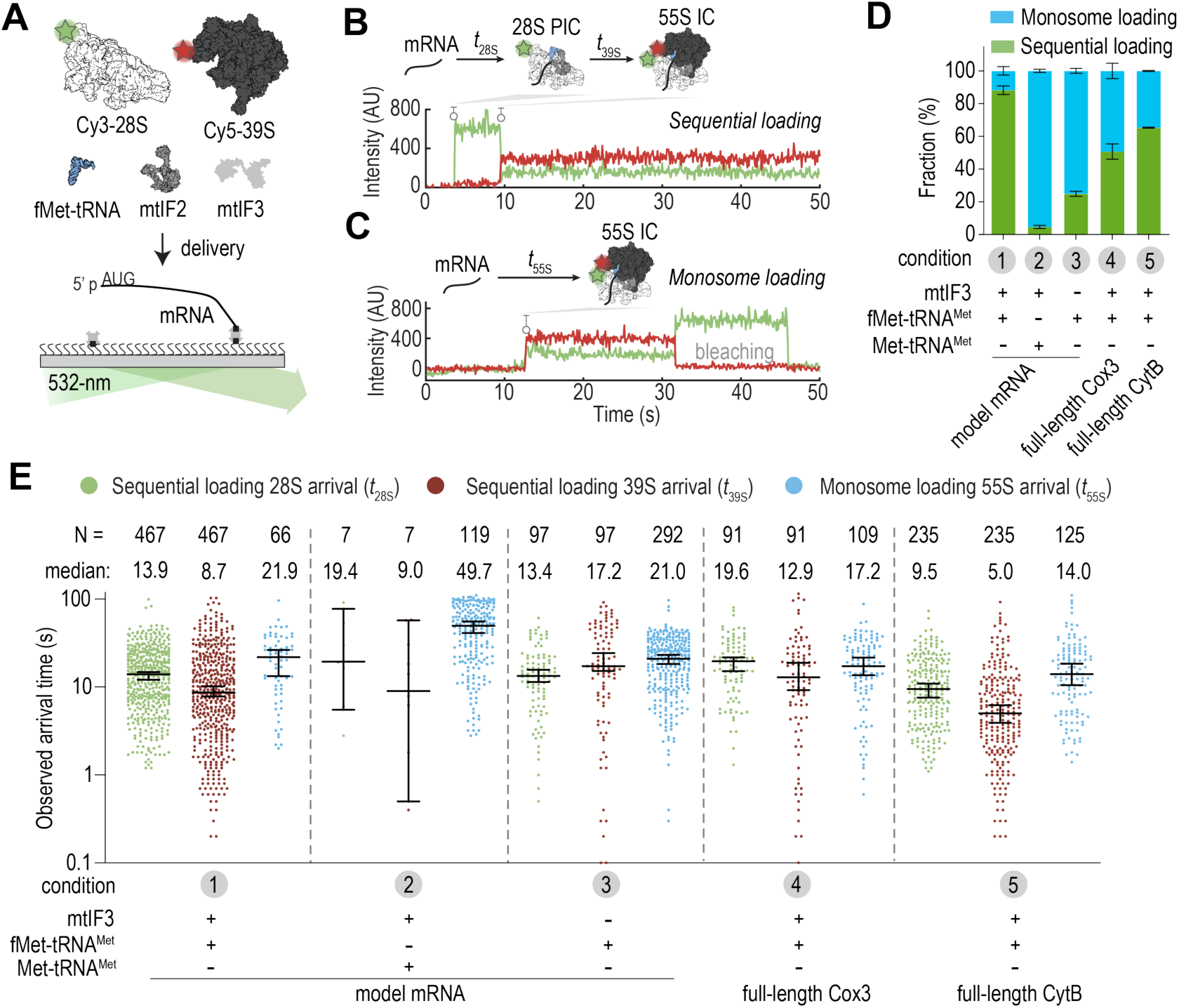
Two parallel pathways of leaderless mRNA initiation. **(A)** Schematic of assay to track Cy3-28S and Cy5-39S binding to tethered leaderless mRNAs. **(B)** Example fluorescence trace showing sequential loading of the 28S (Cy3, green) and 39S (Cy5, red) to the mRNA. **(C)** Example fluorescence trace showing simultaneous binding of the 28S (Cy3, green) and 39S (Cy5, red) to the mRNA as a monosome. **(D)** Frequencies for sequential loading and monosome loading pathways under different reaction conditions. Indicated errors are standard error of the mean from two biological replicates. **(E)** Observed 28S, 39S, or 55S monosome arrival times in experiments under different conditions. Each point represents a binding event. The numbers of observed molecules (N) and median values are listed on top of the plots, and errors bars are 95% confidence intervals.

Notably, when the Met-tRNA^Met^ was not *N*-formylated, only ∼5% of the initiation events on the model mRNA occurred via the sequential loading pathway (Figure 5D). This observation was consistent with the impaired 28S PIC formation with Met-tRNA^Met^ (Figures S5E-S5G), and suggested that the monosome loading pathway can initiate with Met-tRNA^Met^, albeit at a ∼2.5-fold slower rate (Figure 5E). When mtIF3 was omitted, the sequential loading pathway frequency also substantially decreased (to ∼25%, Figures 5D and 5E). This, however, was not due to disrupted 28S PIC formation (Figure S4D), but rather plausibly due to increased monosome formation in the absence of mtIF3 prior to mRNA binding^32^. Together, these results suggested that the initiation pathways can be dynamically regulated.

### Both pathways yield elongation-competent 55S complexes

We next wondered whether the 55S complexes assembled from both pathways were active in accepting an elongator tRNA to decode the second codon displayed in the ribosomal A site. To the tethered leaderless mRNAs that contained a phenylalanine UUC codon as the second codon, a delivery mixture with 1.5 nM Cy3-28S, 4 nM Cy5-39S, 20 nM Cy5.5-mtIF2, 200 nM fMet-tRNA^Met^, 300 nM mtIF3 and 20 nM mtEF-Tu•GTP•Phe-tRNA^Phe^ ternary complex (Cy3.5-Phe-TC, dye labeled on the tRNA^Phe^) was added to start the reaction. Real-time data acquisition was simultaneously started with dual illumination by the 532- and 640-nm lasers to directly excite all the four fluorescently labeled components (Figure 6A).

**Figure 6.**
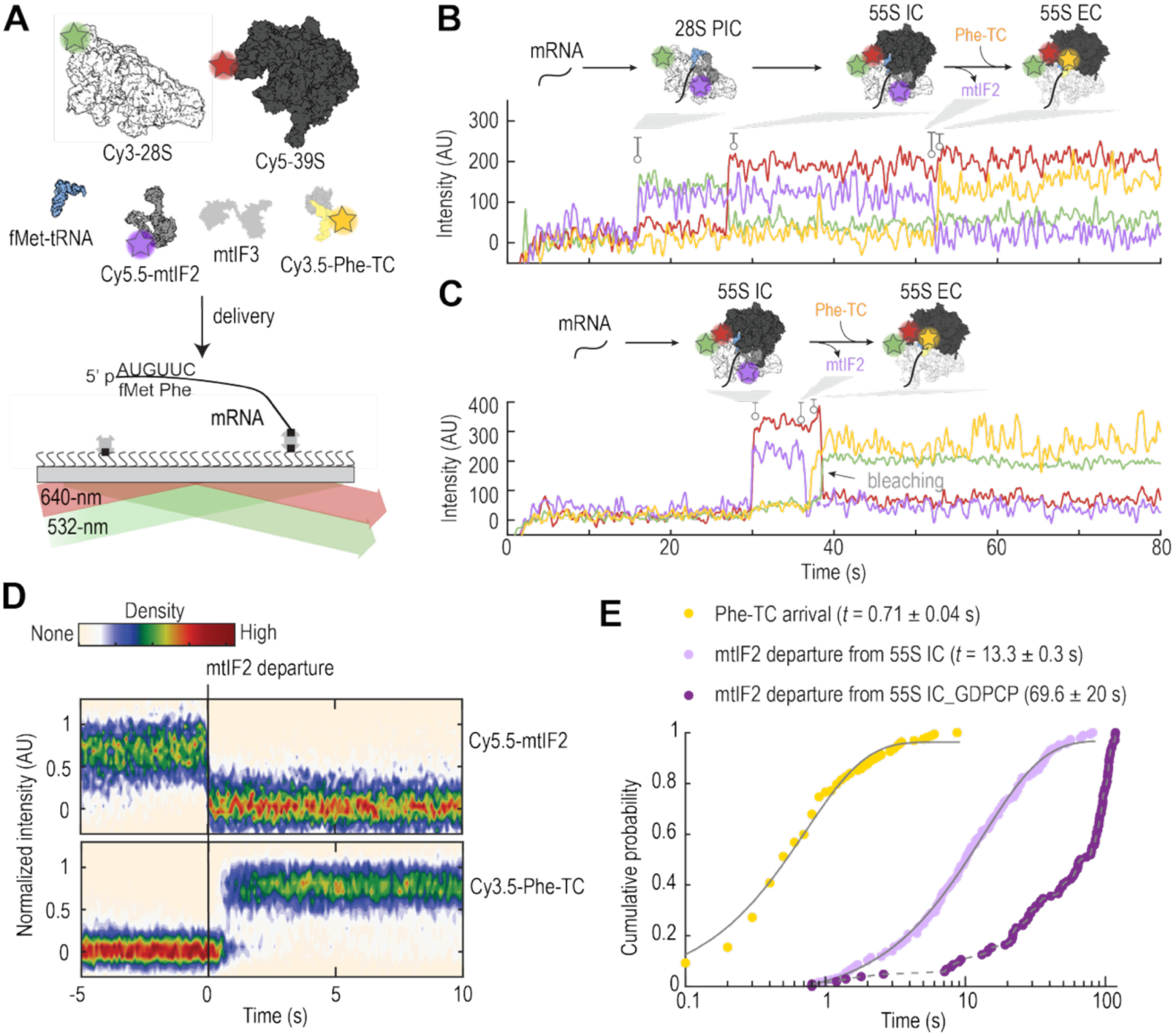
The transition from initiation to elongation is gated by mtIF2. **(A)** Schematic of assay to track Cy3-28S (green), Cy5-39S (red), Cy5.5-mtIF2 (purple), and mtEF-Tu•GTP•Phe-(Cy3.5)-tRNA^Phe^ (Cy3.5-Phe-TC, yellow) binding to tethered leaderless mRNAs using dual illumination with the 532- and 640-nm lasers. Example fluorescence trace showing direct tracking of 28S, 39S, mtIF2, and Phe-TC in the sequential loading **(B)** and monosome loading **(C)** initiation pathways. **(D)** Density heat map of the normalized fluorescence intensities of Cy5.5-mtIF2 and Cy3.5-Phe-TC synchronized to the departure of mtIF2 from the ribosomal complex (N = 186). **(E)** Observed lifetime of mtIF2 on 55S IC (light purple) and the subsequent Phe-TC arrival time (yellow) in the presence of GTP (N = 186). The distributions were fit to a single-exponential model and mean times (± 95% CI) are shown. The observed lifetime of mtIF2 on 55S IC in the presence of GDPCP is shown in dark purple (median lifetime ± 95% CI, N = 104).

As expected, initiation via the two parallel pathways were also observed (Figures 6B and 6C), and more importantly, we observed Cy3.5-Phe-TC binding to the 55S IC assembled from both pathways. Correlating the Cy5.5-mtIF2 signals to ribosome signals revealed that mtIF2 co-arrived with the 28S subunit (Figure S9A) or the 55S monosome (Figure S9B) to bind the mRNA. Omitting either mtIF2 or fMet-tRNA^Met^ suppressed ribosome recruitment (Figure S9C). After 55S IC assembly on the mRNA, mtIF2 resided on this complex for ∼13 s before its departure (dissociation rate ∼0.08 s^-1^), which was followed by rapid binding of the Cy3.5-Phe-TC (∼1.4 s^-1^) (Figures 6B-6E). The dissociation of mtIF2 required GTP hydrolysis by this factor. When the non-hydrolysable GDPCP analog was used, the occupancy time of mtIF2 on the 55S IC was extended by ∼6-fold (median lifetime ∼70 s, and likely photobleaching limited) and the binding of Cy3.5-Phe-TC was inhibited (Figures 6E and S9D). Thus, both initiation pathways produce active complexes, and their progression to translation elongation is gated by mtIF2.

## Discussion

While prior studies have established the key components of mammalian mitochondrial translation initiation on leaderless mRNAs, a coherent model for this central biological process remain elusive^2,16–20,22,25^. Here, we directly monitored leaderless mRNA translation initiation up to the transition to elongation, revealing its heterogeneous pathways and the kinetics of individual sub-steps. Synthesizing our results with previous reports^2,16–20,22,25^, we propose a quantitative model for leaderless mRNA translation initiation by human mitochondrial ribosomes (Figure 7).

**Figure 7.**
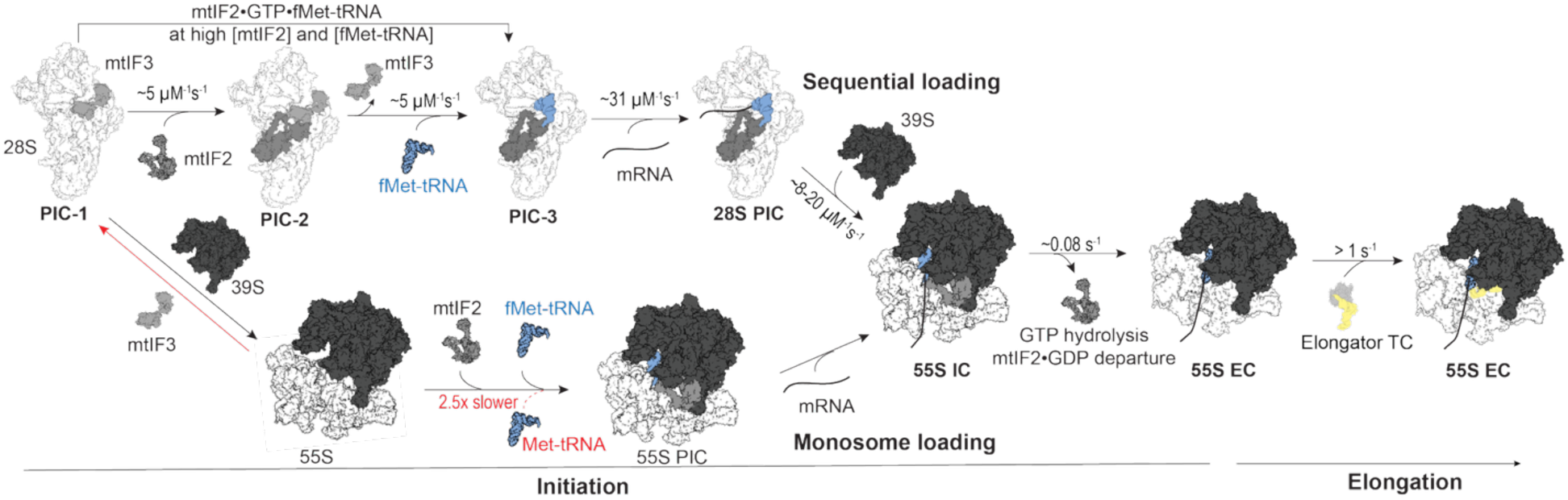
A quantitative model of human mitochondrial leaderless mRNA translation initiation. The mtIF3 protein is bound on the 28S subunit from its late biogenesis stages. Upon 28S subunit maturation, mtIF2 and fMet-tRNA^Met^ can bind to the subunit individually or as a complex. This enables rapid loading of the leaderless mRNA to form a 28S PIC that is poised to 39S joining. Alternatively, the 28S subunit can associate with the 39S to form a 55S monosome. Upon mtIF2 and fMet-tRNA^Met^ binding to the 55S, this complex becomes competent in mRNA binding to initiate. Once a 55S IC is formed, GTP hydrolysis by mtIF2 is activated to promote its release from the ribosome, permitting the first elongator tRNA binding to enter the translation elongation phase. The monosome loading pathway can use non-formylated Met-tRNA^Met^ to initiate at a slower rate, and the usage of this pathway is be limited by mtIF3.

Initiation can occur on the 28S subunits, as demonstrated by our single-molecule observations (Figures 1-3) and the cryo-EM structure of the 28S PIC (Figure 4). The mtIF2 and fMet-tRNA^Met^ bind either sequentially or as a complex to the 28S subunit (PIC-3), from which mtIF3 has dissociated to vacant the binding site on the 28S subunit for fMet-tRNA^Met^ binding^17^. Next, the leaderless mRNA is recruited to bind the PIC-3 (Figures 3F and 3G), forming a 28S PIC (Figure 4) that is poised for 39S subunit joining to form a 55S IC^18,20^. In parallel, the individual mitochondrial ribosomal subunits can also preassemble a 55S monosome and load onto a leaderless mRNA (Figure 7)^16,17^. To enable this pathway, the 55S monosome also needs to be pre-loaded with mtIF2 (Figure S9B) and fMet-tRNA^Met^ (Figure S9C) to promote mRNA binding.

In both initiation pathways, the initial mRNA binding on the 28S subunits might occur at the mS39 region, which is a pentatricopeptide repeat domain-containing protein with mRNA binding activities^22,33^. Upon initial binding, the mRNA may be rapidly (faster than our imaging frame rates) threaded through the uS5m latch towards the P site (Figure S1)^3,4^. The threading might be driven by the mRNA binding energy, which however is insufficient to propagate a long 5’ UTR beyond the P site to enable initiation at an internal start site (Figures S4A, S4B, and S4F)^18^. Once the 5’ start codon reaches to the P site and is recognized by the fMet-tRNA^Met^, the mRNA is clamped on the ribosome to commit to translation.

While both initiation pathways yield 55S initiation complexes that are functional in entering the polypeptide elongation phase (Figure 6), the relative pathway efficiencies may under tight regulation. In mammalian mitochondria, the *N*-formylation of Met-tRNA^Met^ by the mitochondrial methionyl-tRNA formyltransferase (MTFMT)^34^ plays an important role in partitioning the single tRNA^Met^ into translation initiation and elongation. Our results revealed that the monosome loading pathway is however less stringent in discriminating Met-tRNA^Met^ (Figures 5D and 5E), permitting slower initiation with Met-tRNA^Met^ when MTFMT activity is compromised. This initiation pathway could plausibly account for the basal mitochondrial translation activities that are maintained in MTFMT knock-out cells^35^ or in cells derived from Leigh syndrome patients with MTFMT mutations^36,37^.

A potential mechanism to limit the usage of the promiscuous monosome loading pathway is exerted by mtIF3. The mtIF3 protein is already bound on the 28S subunit from its late biogenesis stages (PIC-1, Figure 7)^5,17,38^, providing a rapid pathway for the newly assembled 28S subunit to recruit mtIF2 and fMet-tRNA^Met^ and initiate on leaderless mRNAs (Figures 2, 3, and 4). The mtIF3 can also prevent the formation of a 55S monosome devoid of mRNAs^32^, e.g. after ribosome recycling^39^, to further suppress the monosome loading pathway (Figure 5D).

Finally, the transition from translation initiation to elongation is controlled by mtIF2 GTP hydrolysis and release from the ribosome^22^ (Figures 6A-6E). The occupancy time of mtIF2 on the 55S IC is ∼13 s (Figures 6E and 7), similar to its counterpart in yeast cytosolic translation initiation^40^, wherein the homologous eIF5B protein resides on the 80S IC for a similar duration to kinetically proofread the start site selection and the identity of the amino acid carried by the initiator tRNA^41,42^. Whether mtIF2 plays a similar role in safeguarding the fidelity of leaderless mRNA translation initiation warrants in-depth examinations.

Collectively, our results demonstrate that human mitochondrial leaderless mRNA translation initiation can occur through multiple parallel pathways (Figure 7), underscoring the heterogeneous nature of translation initiation in all studied systems^7,30,43,44^. The reconstituted human mitochondrial translation system, the single-molecule approaches, and the quantitative framework we developed here lay the foundation for future endeavors towards a dynamic understanding of human mitochondrial translation and its regulation in health and disease^2^.

## Acknowledgements

We thank Nenad Ban and Tanja Schönhut for the mtIF2, mtIF3, MTFMT and MetRS overexpression plasmids; Shintaro Iwasaki and Hironori Saito for a detailed on-gel mito-FUNCAT assay protocol; Chuo Chen and Yong Lu for the purification of CoA-dye conjugates; Letian Bao and Anthony C. Forster for suggestions to improve tRNA charging efficiencies; Zheng Xing and Benjamin P. Tu for characterizing the fMet-tRNA using mass spectrometry; Kenneth D. Westover lab for sharing equipment; Joshua T. Mendell, Israel S. Fernández, Joseph D. Puglisi and the Puglisi lab members for discussion. We thank Andy Lemoff and the UT Southwestern Proteomics Core facility for assistance with proteomics analysis of the purified mitochondrial ribosome samples. We thank Angela Mobley and the Flow Cytometry Core Facility at UT Southwestern Medical Center for assistance with flow cytometry. We thank Yan Han, Yang Li, Joyce Fung and James Chen at the Structural Biology Lab (SBL) at UT Southwestern Medical Center for cryo-EM sample preparation and data collection. The SBL is partially supported by grant RP220582 from the Cancer Prevention & Research Institute of Texas (CPRIT). All cryo-EM data was collected at the UT Southwestern Cryo-Electron Microscopy Facility (CEMF). The CEMF is supported by a core facilities award from CPRIT (RP220582). This work was supported by the Welch Foundation (I-2218-20240404 to J.W. and I-2246-20250403 to D.L.K.), NIGMS (R00GM141261 and R01GM155152 to D.L.K.), and the UTSW Endowed Scholars Fund (D.L.K. and J.W.).

## Author contributions

J.W. conceived the project. S.S. established the edited cell lines, performed the on-gel mito-FUNCAT assays, and purified the mitochondrial ribosomal subunits. Y.X. prepared the protein factors. J.W. performed and analyzed all single-molecule experiments. J.W. prepared the cryo-EM samples, and D.L.K analyzed cryo-EM data and modeled the structure with input from J.W. The manuscript was written by J.W. with input from all authors. All authors read and approved the final manuscript.

## EXPERIMENTAL MODEL AND SUBJECT DETAILS

HEK293S GnTI^-^ cells were obtained from ATCC (CRL-3022). To guard against genomic instability, an aliquot of the cell line was passaged for only 4−6 weeks. Cells were confirmed to be free of mycoplasma using InVivogen MycoStrip kit (#rep-mys-10). Cell lines were not validated further. Cells were thawed in Dulbecco’s modified Eagle’s medium (DMEM) (high glucose) medium (Gibco #11965118) supplemented with 10% v/v fetal bovine serum (FBS, GeminiBio #900-108-500), 100 units/ml penicillin and streptomycin (Gibco #15150-122), and 2 mM L-glutamine (Gibco #25030-081), and passaged in monolayer culture at 37 °C in 5% CO_2_. HEK293S GnTI^-^ cell lines with mitoribosomal protein tagged were created via CRISPR-Cas9 editing (see below). For suspension culture of wild type and edited HEK293S cells, after expanding to eight 15 cm dishes, cells were sloughed off and transferred to suspension culture in FreeStyle 293 medium (Gibco #12338-018) supplemented with 2% v/v FBS, 100 units/mL penicillin and streptomycin. Suspension cells were grown with orbital shaking at 130 rpm in baffled flasks at 37 °C in 8% CO_2_. Cells were passaged to maintain a suspension density within the range of 0.4 × 10^6^ cells/ml to 3 × 10^6^ cells/mL.

## METHOD DETAILS

### CRISPR-Cas9 & Homology-directed repair

#### Sequences

To generate the mS26-A1, mS31-S6 and bL31m-S6 cell lines, the guide sequences GGACTCCTAGGGGCCCAGTA (mS26), ATACAGTTCAATTAAGACCA (mS31), and TCCCGGGGTGGCTGGAGCCA (bL31m) were cloned into the pX458 vector backbone using the published approach^55^. To insert the tags onto the endogenous copies of the genes, single-stranded DNA ultramer repair templates were purchased from Integrated DNA Technologies that contained about 40-60 nucleotides of flanking sequence on either side of the desired insertion. See Table S1 for all guide oligo, repair template, and PCR screening oligo sequences.

#### Transfections, sorting, & screening overview

Approximately 24 hours post seeding in a well of a 6-well plate, wild-type HEK293S cells at ∼60% confluency were transiently transfected (Liopfectamine 3000, ThermoFisher #L13000008) with 1 µg of the relevant pX458 plasmid and 2 µg of ssDNA repair template. Cells were allowed to recover for 48 hours. Single, eGFP-positive cells were sorted at the UT Southwestern Flow Cytometry Core Facility into a well of a 96-well plate that contained 50% conditioned DMEM (high glucose) medium. Individual colonies were allowed to recover until they were visible by eye (approximately 2 weeks), upon which colonies were transferred to a well of a 24-well plate. Once confluent, colonies were screened via PCR, Sanger sequencing, and Western blot analyses. Successfully edited cell lines were expanded and ultimately stored as stocks in 10% DMSO/ 90% FBS solution.

#### PCR screening & Sanger sequencing

Genomic DNA was isolated from candidate cell lines using the QuickExtract DNA Extraction Solution (Lucigen, #QE09050) essentially as described, except 10 µL of resuspended cells were added to 50 µL of QuickExtract solution. Following extraction, 1 µL of genomic DNA was added to a standard 24 µL 2X GoTaq Green PCR (Promega, #M7123), typically ∼35 cycles. Sequences for screening oligos are available in Table S1. Following amplification, PCR reactions were analyzed using 2% agarose gel electrophoresis. PCR products with the desired insertion size were submitted for Sanger sequencing, following their purification.

#### Western blots

Whole cell lysates were analyzed via SDS-PAGE followed by transfer to PVDF membranes. All antibodies were diluted in EveryBlot Blocking buffer (BioRad, # 12010020). The following primary antibodies were used: anti-mS26 at 1:2000 from Abcam (ab181863); anti-mS31 at 1:1000 from Thermo Scientific (PA5-109985); and, anti-bL31m at 1:2000 from Thermo Scientific (PA5-97981). Incubations with primary antibodies were for 16 hours at 4 °C. The secondary antibody was Mouse anti-rabbit IgG-HRP (sc-2357) and was used at 1:2,000. Blots were imaged via chemiluminescence.

### On-gel mito-FUNCAT assays

The on-gel mito-FUNCAT assays were performed as described^27^. Wild type or edited HEK293S cells were seeded at a density of 2.0 × 10^6^ cells per well in 2 mL of DMEM supplemented with 10% FBS, 100 units/ml penicillin and streptomycin (Gibco #15150-122), and 2 mM L-glutamine (Gibco #25030-081) in 6-well plates. The cells were incubated at 37 °C in 5% CO_2_ for 20 hours. After washing the cells with PBS, the cells were incubated in methionine-free DMEM (Gibco #21013-024) supplemented with 100 µg/mL anisomycin (ThermoFisher, #502579290) for 15 min at 37 °C in 5% CO_2_. For the negative controls, to inhibit mitochondrial translation, additional 100 µg/mL chloramphenicol (ThermoFisher, #AC227920250) was included in the medium during incubation. Next, L-homopropargylglycine (HPG; Lumiprobe, #3313-100mg) was added to the culture at final 50 µM, and the cells were further incubated at 37 °C in 5% CO_2_ for 2.5 hours.

After the incubation, cells were washed with ice-cold PBS, detached from the plate using a cell scraper, and transferred to 1.5 mL microcentrifuge tubes. Cells were pelleted by centrifugation at 1,000 × g for 3 min at 4 °C. The cell pellet was resuspended in 50 µL lysis buffer (20 mM Tris-HCl pH 7.5, 150 mM NaCl, 5 mM MgCl_2_, 1% Triton X-100, and protease inhibitors) and incubated on ice for 10 min. The lysate was clarified by centrifugation at 15,000 × g for 20 min at 4 °C. The resulting clarified lysate was transferred to a new 1.5 mL tube for click chemistry reactions.

The click chemistry reaction was performed by incubating the clarified lysate with 30 µM Cy5-azide (Lumiprobe, #A3330), a premixed solution of 9.1 mM CuSO_4_ (Fisher Scientific #AA1417836) and 45.45 mM THPTA (Vector Laboratories #CCT1010100), 100 mM sodium ascorbate (Fisher Scientific #AAA1775922), and 100 mM sodium phosphate buffer at 25 °C for 60 minutes. The reaction was quenched by final 1X Laemmli Sample buffer (BioRad #1610747), and samples were heated at 50°C for 5 min before analyzed by 20% SDS-PAGE.

### Preparation of human mitochondrial ribosomal subunits

The mitochondrial ribosomal subunits were purified essentially as described^24,56^, with some modifications.

#### Purification of mitochondria from HEK293S cells

For each purification, 6 L of wild type or edited HEK293S cell cultures were harvest at a density ∼3 × 10^6^ cells/mL by centrifugation at 1000 rcf for 10 min at 4°C. The cells were washed in ice cold PBS and pelleted by centrifugation at 1000 rcf for 10 min at 4°C. Typically, ∼60 g of cell mass could be obtained. The cells were resuspended in 180 mL cold MIB buffer (25 mM Hepes-KOH pH 7.5, 50 mM KCl, 20 mM Mg(OAc)_2_, 2 mM DTT, protease inhibitors) and gently stirred for 20 min at 4 °C. The cell suspension was transferred to a pre-cooled nitrogen cavitation chamber (Parr Instrument Company #4635) on ice, and then 83 mL cold SM4 buffer (25 mM Hepes-KOH pH 7.5, 50 mM KCl, 20 mM Mg(OAc)_2_, 2 mM DTT, 281 mM sucrose, 844 mM mannitol, protease inhibitors) was added. Next, the nitrogen cavitation chamber was filled with nitrogen gas until the pressure reached 500 psi and kept on ice for 20 min. Upon pressure release from the cavitation chamber, the lysate was collected and centrifuged at 800 x g for 15 min at 4 °C. The supernatant was passed through the cheesecloth (Millipore Sigma #475855-1R) into a beaker kept on ice. To the pellet, 90 mL of MIBSM buffer (3:1 MIB:SM4) was added. The resuspended mixture was homogenized using glass Dounce homogenizer. The homogenate was centrifuged at 800 x g for 15 min at 4 °C and the supernatant was passed through the cheesecloth into a beaker kept on ice. The clarified supernatant fractions from the above two centrifugation steps were combined and centrifuged at 10,000 x g for 15 min at 4 °C. The tight pellet was resuspended in 10 mL MIBSM buffer, and 200 units of Turbo DNase (ThermoFisher #AM2238) was added. The resuspension was rotated on a roller at 4 °C for 20 min. Collect the crude mitochondria by centrifugation at 10,000 x g for 15 min at 4 °C, and a typical yield was ∼2 g wet weight.

#### Purification of mitochondrial ribosomal subunits

The mitochondria pellet was resuspended in 5 mL lysis buffer (25 mM Hepes-KOH pH 7.5, 50 mM KCl, 20 mM Mg(OAc)_2_, 2% Triton X-100, protease inhibitors, RNase inhibitors) and kept on ice for 30 min, then homogenized using glass Dounce homogenizer. The homogenate was kept on ice for another 10 min before centrifugation at 30,000 x g for 20 min at 4 °C. The supernatant was further centrifuged at 105,000 x g for 17 hours at 4 °C in a type 90 Ti fixed-angle rotor. The pellet that contained crude mitochondrial ribosomes was resuspended in dissociation buffer (50 mM Hepes-KOH pH 7.5, 300 mM KCl, 5 mM MgCl_2_, 1 mM DTT, protease inhibitors) and centrifuged at 15,000 x g for 5 min. The clarified solution was mixed with final 1 mM puromycin and incubated at 30 °C for 30 min. The resulting solution was loaded onto a 10-30% sucrose gradient (in dissociation buffer) and centrifuged at 75,416 x g for 18 hours in a SW41 rotor. The gradient was fractionated on a BioComp piston gradient fractionator. The fractions corresponding to 28S and 39S subunits were separately concentrated in a 30 kD MWCO Amicon Ultra centrifugal filter and buffer exchanged into mitoribosome storage buffer (50 mM Tris-HCl pH 7.5, 1 mM DTT, 30 mM KCl, 10 mM MgCl_2_, 6% w/v sucrose, 0.1 mM Spermine, and 1 mM Spermidine) before flash frozen on liquid nitrogen, and stored at -80 °C. The concentrations of mitoribosomal subunits were measured by A_260nm_ using Nanodrop and determined using the conversion: 1 A_260nm_ = 77 pmol 28S and 1 A_260nm_ = 55 pmol 39S.

#### Purification of labeled mitochondrial ribosomal subunits

For labelling of the 28S or 39S subunits through the A1 or S6 tags^26^, a 200-μl labelling reaction was prepared by mixing 100 μL ribosomal subunits, 5 μM AcpS enzyme (for A1-tagged subunits) or 2 μM SFP synthase (for S6-tagged subunits) and 10 μM biotin-, Cy3-, or Cy5-CoA conjugates in a buffer containing 50 mM HEPES-KOH pH 7.3, 1 mM DTT and 15 mM MgCl_2_ and 50 mM KOAc. The reaction was incubated at 30 °C for 1 hour. To purify the labelled ribosomes from free dye or biotin, the 200 μl reaction mixture was loaded on top of a 700-μl sucrose cushion (30 mM HEPES-KOH pH 7.3, 1 mM DTT, 50 mM KOAc, 15 mM Mg(OAc)_2_ and 15% w/v sucrose) for ultracentrifugation at 470,000 x g at 4 °C for 90 min in an MLA-130 rotor. The pellet containing labelled mitoribosomal subunits was washed once with and then resuspended in the mitoribosome storage buffer and stored at −80 °C after flash-freezing with liquid nitrogen.

### Purification of unlabeled and labeled mtIF2

The pET24a plasmid for the expression of mature human mtIF2 protein with a N-terminal His_6_-tag and a TEV cleavage site was a kind gift from the Ban lab^22^. For fluorescent labeling of mtIF2, a ybbR-tag (DSLEFIASKLA) was inserted to the plasmid, such that the recombinant protein was expressed as His_6_-TEV site-GGSGSG-ybbR tag-mtIF2. Plasmids were transformed individually into *Escherichia coli* Rosetta2 (DE3) pLysS (Novagen #71403-M), and overexpressed separately in LB media by induction at an OD_600nm_ of 0.6-0.8 with 0.15 mM IPTG at 18 °C. For each purification, 3 L of cells were harvested after 16 hours by centrifugation at 5,000 x *g* for 15 min at 4 °C in a Fiberlite F9 rotor (ThermoFisher), and resuspended in lysis buffer (200 mM KCl, 5 mM MgCl_2_, 50 mM HEPES, 10 mM Imidazole, 10% Glycerol, 5 mM β-mercaptoethanol, pH 7.3 at room temperature) supplemented with 1 mM PMSF, 0.6 µM Leupeptin, 2 µM Pepstatin A, 2 mM Benzamidine. Cells were lysed by sonication (Fisherbrand model 505, 2 s on / 2 s off, 6 min total on time at 40% amplitude), and lysates were cleared by centrifugation at 38,000 x *g* for 30 min at 4 °C in an Fiberlite F21 rotor followed by filtration through a 0.22 μm syringe filter. Clarified lysate was loaded onto a Ni-NTA gravity flow column equilibrated in lysis buffer, washed with 15 column volumes (CV) of lysis buffer, 15 CV of wash buffer (same as lysis buffer but with 1000 mM KCl and 20 mM Imidazole), and 2 CV of lysis buffer. Recombinant proteins were eluted with elution buffer (same as lysis buffer but with 50 mM KCl and 250 mM Imidazole). Fractions with recombinant protein were diluted 1:10 with Low salt buffer (50 mM KCl, 5 mM MgCl_2_, 50 mM HEPES, 10% Glycerol, 5 mM β-mercaptoethanol, pH 7.3 at room temperature) and loaded onto a 5 mL Heparin column (Cytiva #17040601)). After a further 5 CV wash with 9:1 Low salt buffer:High salt buffer (same as Low salt buffer but with 800 mM KCl), proteins were eluted as 1 mL fractions using an 80 mL linear gradient to final 100% High salt buffer. Fractions with recombinant protein were identified by SDS-PAGE analysis and concentrated using a 10 kD MWCO Amicon Ultra centrifugal filter. For unlabeled proteins, the proteins were incubated with TEV protease at RT for 2 hours. The TEV protease and cleaved His_6_ tag were removed via a subtractive Ni-NTA gravity column equilibrated in TEV Buffer (200 mM KCl, 5 mM MgCl_2_, 50 mM HEPES, 10 mM Immidazole, 1 mM DTT, pH 7.3 at room temperature) with the flow-through collected. Unlabeled mtIF2 proteins were subjected to a final purification step using size exclusion chromatography (SEC) on a Superdex 200 column (Cytiva #28990944) equilibrated in SEC buffer (200 mM KOAc, 5 mM MgCl_2_, 40 mM HEPES, 10% glycerol, 1 mM DTT, pH 7.3 at room temperature). For fluorescently labeling of mtIF2, 15 µM His_6_-TEV site-GGSGSG-ybbR tag-mtIF2 was incubated at 30 °C for 30 min with 2 µM Sfp synthase enzyme and 25 µM of either Cy5-CoA, or Cy5.5-CoA substrates, respectively. Then TEV protease was added to this labeling reaction and further incubated at RT for 2 hours. TEV protease, Sfp synthase, and the cleaved His_6_ tag were removed via a subtractive Ni-NTA gravity column equilibrated in TEV Buffer, with the flow-through collected. Free dye was removed via purification over 10DG-desalting columns (Bio-Rad, # 7322010) equilibrated in TEV Buffer. Dye-labeled mtIF2 proteins were subjected to a final SEC purification as for the unlabeled proteins. Fractions containing mtIF2 were concentrated using a 10 kD MWCO Amicon Ultra centrifugal filter (Millipore Sigma #UFC801024), aliquoted, flash frozen on liquid nitrogen, and stored at -80 °C. Protein concentrations were determined via absorption at 280 nm for total protein, and Cy5 or Cy5.5 concentrations for labeled proteins using a nanodrop, respectively. The dye labeling efficiencies were ∼70%.

### Purification of mtIF3

The pET21c-IF3mt plasmid for overexpression of the mature form of human mtIF3 protein was a gift from Linda Spremulli (Addgene plasmid #31078)^57^. The final protein contains an uncleavable His_6_-tag at the C-terminus, and the mtIF3-His_6_ protein is referred to as mtIF3 protein throughout the manuscript. The plasmid was transformed into *Escherichia coli* Rosetta2 (DE3) pLysS and overexpressed in LB media by induction at an OD_600nm_ of 0.6 with 0.5 mM IPTG at 30 °C. For each purification, 5 L of cells were harvested after 5 hours by centrifugation at 5,000 x *g* for 15 min at 4 °C in a Fiberlite F9 rotor, and resuspended in lysis buffer (200 mM KCl, 5 mM MgCl_2_, 50 mM HEPES, 10 mM Imidazole, 10% Glycerol, 5 mM β-mercaptoethanol, pH 7.3 at room temperature) supplemented with 1 mM PMSF, 0.6 µM Leupeptin, 2 µM Pepstatin A, 2 mM Benzamidine. Cells were lysed by sonication (2 s on / 2 s off, 6 min total on time at 40% amplitude), and lysates were cleared twice by centrifugation at 38,000 x *g* for 20 min at 4 °C in an Fiberlite F21 rotor followed by filtration through a 0.22 μm syringe filter. Clarified lysate was loaded onto a Ni-NTA gravity flow column equilibrated in lysis buffer, washed with 15 CV of lysis buffer, 15 CV of wash buffer (same as lysis buffer but with 1000 mM KCl and 20 mM Imidazole), and 2 CV of lysis buffer. Recombinant proteins were eluted with elution buffer (same as lysis buffer but with 100 mM KCl and 250 mM Imidazole). Fractions with recombinant proteins were purified on a 5 mL Heparin column followed by a 5 mL SP column (Cytiva #17115101) in using the same buffers and gradient as described for the purification of mtIF2. The eluted mtIF3-His_6_ protein was subjected to a final SEC purification on a SD200 column (Cytiva #28989336) in the SEC buffer (200 mM KCl, 5 mM MgCl_2_, 40 mM HEPES, 10% Glycerol, 1 mM DTT, pH 7.3). Fractions containing mtIF3-His_6_ were concentrated using a 10 kD MWCO Amicon Ultra centrifugal filter, aliquoted, flash frozen on liquid nitrogen, and stored at -80 °C. Protein concentrations were determined via absorption at 280 nm using Nanodrop.

### Purification of bovine mtEF-Tu

The mature form of bovine mtEF-Tu, which shares 95.8% identity or 97.8% similarity to the mature human mtEF-Tu sequence, was used here in the elongation assay. The pET24(+)-mtEFTu plasmid for overexpression of the protein was a gift from Linda Spremulli (Addgene plasmid #30535)^58^. The final protein contains an uncleavable His_6_-tag at the C-terminus. The plasmid was transformed into *Escherichia coli* Rosetta2 (DE3) pLysS and overexpressed in LB media by induction at an OD_600nm_ of 0.6 with 0.15 mM IPTG at 28 °C. For each purification, 3 L of cells were harvested after 4.5 hours by centrifugation at 5,000 x *g* for 15 min at 4 °C in a Fiberlite F9 rotor, and resuspended in lysis buffer (300 mM KCl, 5 mM MgCl_2_, 20 mM HEPES, 10 mM Imidazole, 10% Glycerol, 5 mM β-mercaptoethanol, pH 7.3 at room temperature) supplemented with 1 mM PMSF, 0.6 µM Leupeptin, 2 µM Pepstatin A, 2 mM Benzamidine. Cells were lysed by sonication (2 s on / 2 s off, 6 min total on time at 40% amplitude), and lysates were cleared twice by centrifugation at 38,000 x *g* for 20 min at 4 °C in an Fiberlite F21 rotor followed by filtration through a 0.22 μm syringe filter. Clarified lysate was loaded onto a Ni-NTA gravity flow column equilibrated in lysis buffer, washed with 15 CV of lysis buffer, 15 CV of wash buffer (same as lysis buffer but with 350 mM KCl and 20 mM Imidazole), and 2 CV of lysis buffer. Recombinant proteins were eluted with elution buffer (same as lysis buffer but with 100 mM KCl and 250 mM Imidazole). Fractions with recombinant proteins were subjected to a final SEC purification on a SD200 column (Cytiva #28990944) in the SEC buffer (150 mM KOAc, 1 mM MgCl_2_, 30 mM HEPES, 10% Glycerol, 1 mM DTT, pH 7.3). Fractions containing mtEF-Tu were concentrated using a 10 kD MWCO Amicon Ultra centrifugal filter, aliquoted, flash frozen on liquid nitrogen, and stored at -80 °C. Protein concentrations were determined via absorption at 280 nm using Nanodrop.

### Preparation of tRNAs

Due to technical challenges in obtaining purified human mitochondrial tRNAs in sufficient quantities for our in vitro studies, we have used an unmodified human mitochondrial tRNA^Met^ in translation initiation assays^22,59^ and Cy3.5 labeled *E. coli* Phe-tRNA^Phe^ for translation elongation assays^40^.

#### Unlabeled human mitochondrial tRNA^Met^

The tRNA^Met^ was either purchased from TriLink or transcribed from a DNA template with a 5’-end T7 promoter and hammerhead ribozyme. For transcribed *tRNA^Met^,* the plasmid (ordered from Genscript) was linearized via digestion with BstNI for transcriptional termination at the 3’-CCA of tRNA^Met^. The tRNA was transcribed with the MEGAScript T7 Transcription kit (ThermoFisher #AMB13345). Purified and linearized plasmid was used as the template for in vitro transcription with T7 polymerase at 37 °C for 4 hours at 16 mM MgCl_2_, during which the ribozyme self-cleaved (>80% efficiency). Mature tRNA^Met^ was separated from precursor RNA and cleaved ribozyme via 10% acrylamide gel electrophoresis in the presence of 8 M urea. After excision of the tRNA^Met^ band, the RNA was extracted 3x at room temperature for 12 hours with 300 mM ammonium acetate, and ethanol precipitated. tRNA^Met^ was resuspended in 10 mM NaCl,10 mM Bis-tris, pH 7.0 and stored at -80°C.

#### Cy5-labeled human mitochondrial tRNA^Met^

HPLC purified, synthetic human mitochondrial tRNA^Met^ with U17 replaced with Uridine-C6 Amino linker was purchased from TriLink. The modified tRNA^Met^ was labeled with excess sulfo-Cy5-NHS ester (Lumiprobe, #13320) in a buffer containing 50 mM Hepes-KOH, pH 8.0 and 0.9 M NaCl at 30 °C for 2 hours followed by 4 °C for 16 hours in the dark. The labeling reaction was stopped by addition of final 0.3 M NaOAc (pH 5.2). The tRNAs were extracted twice with phenol (pH 4.7) and once with chloroform before precipitated in 3 volumes of ethanol. The precipitated tRNAs were pelleted by centrifugation at 20,000 x g at 4 °C for 1 hour and dissolved in 75 µL ddH_2_O. The resuspended tRNA solution was loaded to a Bio-Spin 6 column (BioRad, # 7326221) pre-equilibrated with ddH_2_O to remove any excess free dyes. The labeling efficiency was ∼100% based on absorbance measurements on a Nanodrop for A_260nm_ for total RNA and Cy5. The purified Cy5-tRNA^Met^ was flash-frozen on liquid nitrogen and stored at -80 °C.

#### Charging and formylation of labeled and unlabeled tRNA^Met^

To charge the unlabeled for Cy5-labeled tRNA^Met^, the tRNAs were heated at 85 °C for 10 min and slowly cooled to RT. Then the tRNAs were added to final 5 µM in a charging reaction buffered with 50 mM Tris-HCl (pH 8.0 at room temperature), 10 mM MgCl_2_, 0.2 mM Spermine, 0.2 mg/mL BSA, 20 mM KCl, and 5 mM DTT. The charging reaction was initiated with the addition of final 2.5 mM ATP, 2 mM L-methionine, and 1.5 µM MetRS. The reaction was incubated at 37 °C for 30 min. If non-formylated Met-tRNAs were desired, the reactions were stopped by adding final 0.3 M NaOAc pH 5.0 and kept on ice until further purification. To formylate the Met-tRNAs, final 1 mM 10-formyltetrahydrofolate and 1 µM MTFMT^22^ was added to the charging reaction mixture and further incubated for 15 min at 37 °C. Upon stopping the reactions with final 0.3 M NaOAc pH 5.0, the tRNAs were extracted once with phenol (pH 4.7) and once with chloroform. The water phase that contained tRNAs was mixed with 3 volumes of ethanol and kept at -20 °C for 30 min before centrifugation at 20,000 x g for 30 min at 4 °C. The tRNA pellet was dissolved in 70 µL tRNA storage buffer (2 mM Mg(OAc)_2_ and 10 mM NaOAc pH 5.2). The charged tRNA was further purified with a Bio-Spin 6 column that were equilibrated with tRNA storage buffer. The charging efficiency was > 90% based on acid urea PAGE analyses of the final tRNA product.

### Preparation of leaderless mRNAs

The chemically synthesized model leaderless mRNAs were purchased from Integrated DNA Technologies (IDT) or Millipore Sigma. To produce full-length human mitochondrial Cox3 and CytB leaderless mRNAs, the MT-COX3 and MT-CYTB genes were individually cloned into a pUC57 vector with a 36-nucleotide polyA sequence at the 3’ end (Genscript). The mRNA gene sequences were located to the 3’ of a T7 promoter sequence and a hammerhead ribozyme sequence^22,59^. The DNA templates for in vitro transcription of the mRNAs were amplified by PCR from the plasmids and purified with a Qiagen PCR purification kit. In vitro transcription with the MEGAScript T7 Transcription kit (ThermoFisher #AMB13345) was performed at 37 °C for 5 hours and transcripts were incubated for 1 hour at 60 °C to complete hammerhead cleavage. The mRNA was separated from the hammerhead ribozyme using 10% acrylamide gel electrophoresis in the presence of 8 M urea. After excision of the appropriate band, the mRNA was extracted over night at 4 °C by shaking the gel pieces in 300 mM ammonium acetate and ethanol precipitated. The mRNA was dissolved in ddH_2_O and stored at -80°C. To label the 3’ end of the full-length Cox3 and CytB mRNAs with a Cy3 dye or biotin, the mRNAs were oxidized with potassium periodate and then reacted with Cy3- or biotin-hydrazide, as described^60^.

### Single-molecule assays

#### Instrumentation

We performed all single-molecule experiments at 30 °C (or otherwise denoted) using a custom-built, objective-based (Nikon CFI Apochromat TIRF 60X Oil) total internal reflection fluorescence (TIRF) microscope (Nikon Ti2E inverted microscope), equipped with a MultiSplit V2 system (Cairn Research), Photometrics Kinetix sCMOS camera (Teledyne Photometrics), a LUN-F 488/532/640nm laser system (maximal output power for the three laser lines are 60/50/40 mW), a Legato 110 syringe pump (KD Scientific), and an H201-enclosure (Okolab) for temperature control. Within the MultiSplit, a dichroic beam splitter separated Cy3/Cy3.5 and Cy5/Cy5.5 emission (T635LPXR), a filter cube (ET667/30m, T685LPXR, ET720/60m, Chroma) was used to separate Cy5 from Cy5.5 emission signals, and another filter cube (ET615/40m, T585LPXR, ET572/23m, Chroma) was used to separate Cy3 from Cy3.5 emission signals. For experiments with a single 532-nm laser excitation, the laser power was used at 30%; for ALEX excitation, the 532-nm laser was set at 30% power while the 640-nm laser at 20%; for simultaneous illumination with 532- and 640-nm lasers, the laser power was 30% and 20% for the two lasers, respectively. Movies were acquired using the Nikon NIS-Elements software package (v 6.02.03).

#### Imaging surface and buffer conditions

Glass TIRF slides and cover slips were passivated and functionalized as described^61^. Prior to imaging, the chamber was washed with TP50 buffer (50 mM Tris-HCl pH 7.5, 50 mM KCl) and coated with neutravidin (Thermo Fisher #31000) by a 5 min incubation with 75 nM neutravidin diluted in TP50 buffer supplemented with 1µM of DNA blocking oligos (Table S1) and 0.7 mg/mL UltraPure BSA (Thermo Fisher #AM2618). Then the chamber was washed with TP50 buffer and 1X Initiation buffer (30 mM KCl, 10 mM MgCl_2_, 0.1 mM Spermine, 1 mM Spermidine, 50 mM Tris-HCl, pH 7.5 at room temperature)^16^. For reaction imaging, the chamber was washed with Imaging buffer [1X Initiation buffer supplemented with 2 mM GTP (or GDPCP when denoted), casein (62.5 µg/mL) and an oxygen scavenging system^62^: 2 mM Trolox (TSY), 2.5 mM protocatechuic acid (PCA), and 0.06 U/µL protocatechuate-3,4-dioxygenase (PCD)].

#### Experimental setup for *Figure 1* and related experiments

Prior to tethering, 2 nM biotin-28S subunits were incubated at 30 °C for 10 min in 1X Initiation buffer. For tethering, 0.4 nM biotin-28S subunits (diluted in 1X Initiation buffer) were incubated on the neutravidin-coated surface for 5 min at room temperature. For the biotin-block control experiment, prior to 28S tethering, the surface was treated with saturating biotin solution for 5 min at room temperature and washed with TP50 buffer. Non-tethered ribosomes were washed away with 1X Initiation buffer followed by Imaging buffer prior to imaging. Upon start of data acquisition at 200 ms frame rate, a mixture was delivered to the surface that contained 1 nM Cy3-mRNA, 400 nM mtIF2, 300 nM mtIF3 and 200 nM fMet-tRNA^Met^, or otherwise as denoted (e.g. with factor dropout). The surface was excited with the 532-nm laser, and the movie length was 4 min.

#### Experimental setup for Figure S5

The biotin-28S subunit tethering was as described above. Upon start of data acquisition at 100 ms frame rate, a mixture was delivered to the surface that contained 1 nM Cy3-mRNA, 20 nM Cy5-mtIF2, 300 nM mtIF3 and 200 nM fMet-tRNA^Met^ (or Met-tRNA^Met^). The surface was excited with the 532-nm laser, and the movie length was 2 min or otherwise as denoted.

#### Experimental setup for *Figure 2* and related experiments

The biotin-28S subunit tethering was as described above. Upon start of data acquisition at 150 ms frame rate, a mixture was delivered to the surface that contained 200 nM fMet-tRNA^Met^, and Cy3-mRNA, Cy5-mtIF2 and mtIF3 at indicated concentrations. The surface was excited using ALEX with the 532- and 640- nm lasers, and the movie length was 2 min.

#### Experimental setup for *Figure 3* and related experiments

The biotin-28S subunit tethering was as described above. Upon start of data acquisition at 150 ms frame rate, a mixture was delivered to the surface that contained Cy5-fMet-tRNA^Met^, Cy3-mRNA, Cy5.5-mtIF2 and mtIF3 at indicated concentrations. The surface was excited using ALEX with the 532- and 640-nm lasers, and the movie length was 2 min.

#### Experimental setup for *Figure 5* and related experiments

Prior to tethering, the 3’ biotinylated mRNAs were diluted in ddH_2_O to 0.1 ∼ 1 µM, heated at 65 °C for 5 min and fast cooled on ice before using. For tethering, 1 nM biotinylated mRNAs (diluted in 1X Initiation buffer) were incubated on the neutravidin-coated surface for 5 min at room temperature. Non-tethered mRNAs were washed away with 1X Initiation buffer followed by Imaging buffer prior to imaging. Upon start of data acquisition at 100 ms frame rate, a mixture was delivered to the surface that contained Cy3-28S, Cy5-39S, mtIF2, fMet-tRNA^Met^ (or Met-tRNA^Met^), and mtIF3 at indicated concentrations. The surface was excited with the 532-nm lasers, and the movie length was 2 min.

#### Experimental setup for *Figure 6* and related experiments

Prior to tethering, the model mRNA with a Phe codon as the second codon and with a 3’ biotin was diluted in ddH_2_O to 1 µM, heated at 65 °C for 5 min and fast cooled on ice before using. For tethering, 1 nM biotinylated mRNAs (diluted in 1X Initiation buffer) were incubated on the neutravidin-coated surface for 5 min at room temperature. Non-tethered mRNAs were washed away with 1X Initiation buffer followed by Imaging buffer prior to imaging. The mtEF-Tu•GTP•Cy3.5-Phe-tRNA^Phe^ ternary complex (Cy3.5-Phe-TC) was prepared by incubating 20 µM mtEF-Tu and 0.4 µM Cy3.5-Phe-tRNA^Phe^ in 1X Initiation buffer supplemented with 2 mM GTP at room temperature for 15 min. Upon start of data acquisition at 100 ms frame rate, a mixture was delivered to the surface that contained 1.5 nM Cy3-28S, 4 nM Cy5-39S, 20 nM Cy5.5-mtIF2, 200 nM fMet-tRNA^Met^, 300 nM mtIF3, and 20 nM (based on tRNA concentration) Cy3.5-Phe-TC. The surface was excited with the 532- and 640-nm lasers simultaneously, and the movie length was 2 min.

## QUANTIFICATION AND STATISTICAL ANALYSIS

### Single-molecule data analysis

Experimental movies that captured fluorescence intensities over time were processed using MATLAB (R2018b or R2024a), and single-molecule traces were extracted using the Spartan software package (v.3.7.0)^51^, as described previously^40,52,53^. In Spartan, the channels for the different wavelengths were aligned according to the alignment parameters obtained with a movie that was acquired with a TetraSpeck beads (Thermo Fisher #T7279) slide. Fluorescence traces exported from Spartan were further analyzed with MATLAB using published scripts^40,52,53^. Spectral bleedthrough between channels were corrected based on parameters pre-determined using individual dye-labeled DNA oligos. Individual binding events were assigned manually based on the appearance and disappearance of the respective fluorescence signals. Experiments were performed with at least two replicates, and unless intractable due to inefficient complex assembly or inhibition, approximately 100∼400 single molecules were analyzed, indicated by single step photobleaching events.

Arrival times of a particular fluorescent component were defined as the time elapsed from the last molecule event (or the delivery of the labeled component if it was the first binding event in a reaction) until a burst of fluorescence for that component. Lifetimes were defined as the duration of the corresponding fluorescence signal. Kinetic parameters were extracted by fitting the cumulative probability distribution of the observed data to single- or a two-step kinetics model as previously described^29^:

the single-exponential model (for single-step reactions): *C*(*t*) = 1 − *e*^−*b*}*t*^ where *t* is time and *b* is the reaction rate;

the two-step reaction kinetics model: 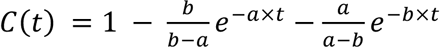 where *t* is time, *a* and *b* are the reaction rates of the two steps.

In concentration titration experiments, the pseudo-first order rate constants were obtained via linear regression model fitting to apparent rate constants in Prism 10 (GraphPad Software Inc.), with reported error ranges denoting 95% CIs. FRET efficiency was defined as *E*_FRET_ = *I*_A_/(*I*_A_+*I*_D_), where *I*_A_ and *I*_D_ represent fluorescence intensities of the acceptor and donor fluorophores. The *E*_FRET_ distributions were fit to single gaussian models, with error ranges denoting the standard deviation of the mean *E*_FRET_. To calculate errors for median association times and lifetimes, bootstrap analyses (n = 10,000) were performed to calculate 95% CI of the observed median using MATLAB (bootstrp and bootci functions).

Statistical details of individual experiments, including number of analyzed molecules (N) and the definition of error bars are indicated in the corresponding figure panels and legends.

### Cryo-EM analysis

#### Sample preparation

A ribosome mix was prepared by mixing 600 nM mtIF2, 800 nM fMet-tRNA^Met^, 300 nM mtIF3, and 300 nM 28S subunits in 1X Initiation buffer supplemented with 2 mM GTP and kept on ice. Separately, the model leaderless mRNA was diluted to 2 µM in the same buffer and kept on ice. In parallel, in-house carbon-coated (3 nm) 300-mesh Quantifoil R2/2 grids (SPI Supplies, #4430C-XA, Lot# 1280410) were glow-discharged for 30 s in a PELCO EasiGlow glow discharger (Ted Pella, conditions: negative charge, 30 mA, 0.38 mBar, 30 s). After grids were prepared, 1.5 µL of the ribosome mix and 1.5 µL of the mRNA mix (both pre-warmed at room temperature) were combined and well mixed at room temperature before being loaded onto the grid. The sample was vitrified by plunging into liquid ethane on a Vitrobot plunger (setup at 4 °C, 100 % humidity, blotting time 3.5 s, wait time 10 s, blotting force 0). The total time elapsed from sample mixing until freezing in liquid ethane was ∼60 s.

#### Cryo-EM data collection

Movies were collected on a Titan Krios equipped with the post-column BioQuantum energy filter (Gatan) and a K3 direct electron detector operated at 300 kV in the super-resolution mode with a pixel size 0.413 Å per pixel using SerialEM software (version 4.0)^46^. The dose rate was 19 e^-^/pixel/s and the total accumulated dose was 50 e^-^/Å^2^ over 50 frames. A nominal defocus range of −1.2 and −2.5 μm was used and the energy filter was set to 20 eV.

#### Cryo-EM image processing

Image processing was done using Cryosparc (v4.5.3)^47^. Imported movies were motion-corrected using Patch Motion Correction and Fourier-cropped to a pixel size of 0.826 Å. Following Patch CTF estimation, a curated set of 8,077 micrographs were selected for subsequent processing. Following rounds of initial manual picking and template picking to generate quality templates from two-dimensional (2D) classes, 1.8 million particles were picked using template picking (Figure S7C). After one round of 2D classification, 521K particles were selected (Figure S7C). Three ab initio maps were generated from 100K particles, and the full dataset was subjected to Heterogenous three-dimensional (3D) Refinement, from which a good 28S subunit class with 296,367 particles was selected (Figure S7D). 293,580 particles were re-extracted in a box of 600 pixels. These particles yielded a map of 3.24 Å with non-uniform 3D refinement^54^, but which suffered from heterogeneity in the density for the tRNA. To sort 28S particles of different compositions, a mask was generated using the structure of mtIF3 when bound to the 28S (PDB 6RW5)^17^ and used for masked 3D classification. 33,600 particles were identified containing density for mtIF3 and mtIF2, and these yielded a 28S–mtIF2–mtIF3 map of 3.01Å (Figures S7H and S7I). Three classes contained strong density for mtIF2 and the fMet-tRNA^Met^; two of these were combined and subjected to further non-uniform 3D refinement and reference-based motion correction for a final set of 107,666 particles and resulted in a 28S PIC map of 2.50Å (Figures S7E and S7F). FSC curves for the 28S-mtIF2-mtIF3 map or 28S PIC map are shown in Figure S7.

#### Model building

PDB 7PO2^5^ was used as the starting model for the 28S PIC structure. The models were manually edited using Coot^48^, ChimeraX^45^, and ISOLDE^49^ and refined using Phenix^50^. Data collection and model statistics are provided in Table S2. ChimeraX was used for analysis of the structure and figure making.

