## Supplementary Materials for "Choreography of human mitochondrial leaderless mRNA translation initiation"

### This PDF file includes:

Figures S1 to S9
Tables S1 to S2

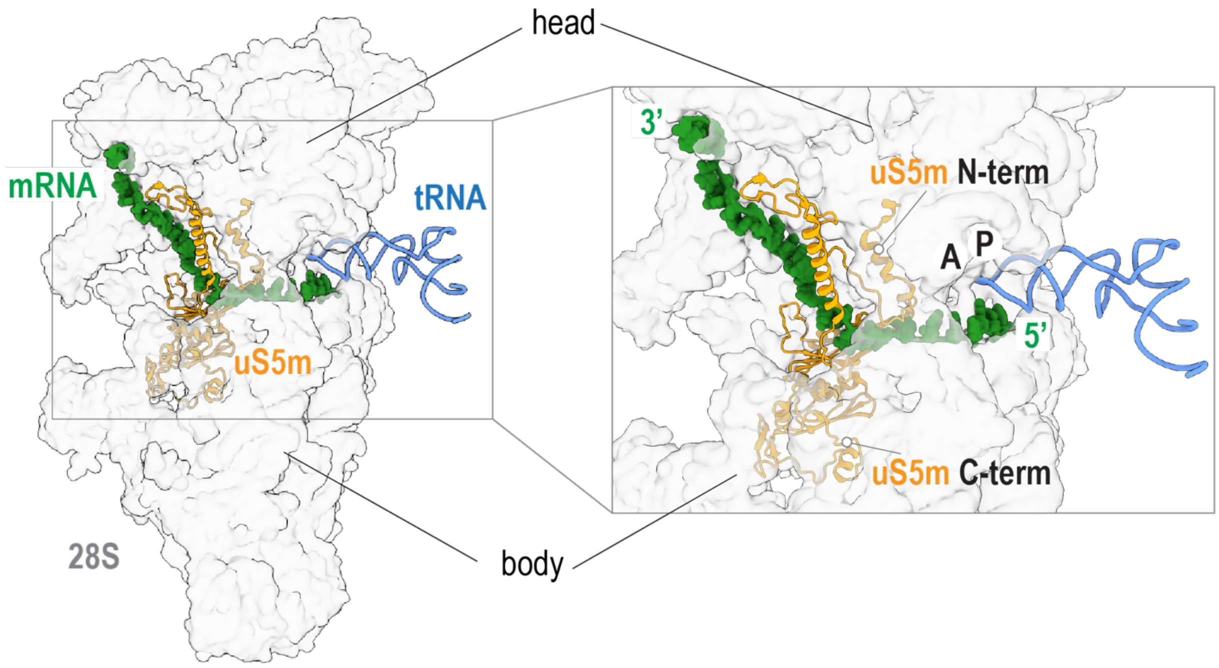

**Figure S1. The mRNA entry path on the 28S subunit.** A structural model (from PDB 7PO2) of the human 28S (gray transparent surface) is shown with the mRNA (green) and P-site tRNA (blue). The N-terminal domain of the ribosomal protein uS5m (orange) is located at the head of the 28S subunit, while the C-terminal domain at the body. These two domains are linked via a helical bridge, connecting the head and the body of the 28S subunit and forming a latch at the mRNA entry path. The ribosomal A- and P-sites and the mRNA 5’ and 3’ ends are indicated in the right panel.

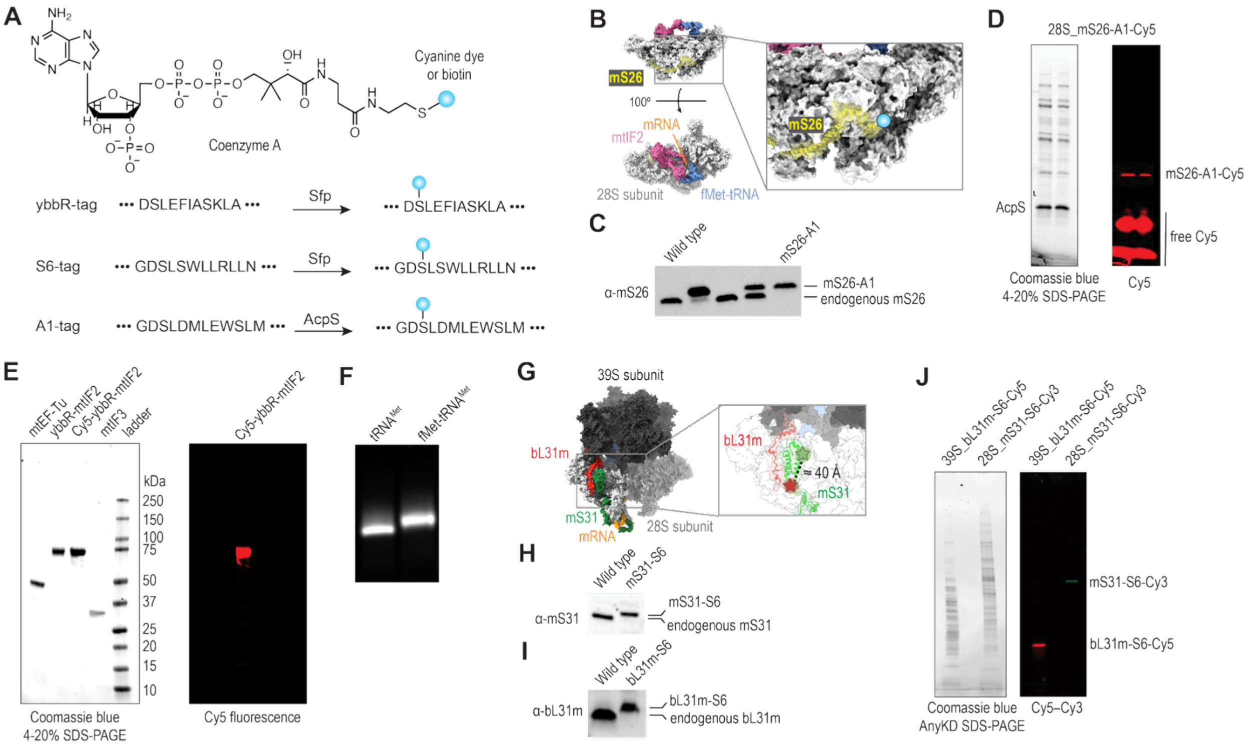

**Figure S2. Reconstitution of human mitochondrial translation initiation. (A)** Peptide tags and the corresponding enzymes that are used for site-specific labeling. **(B)** The C-terminus of mS26 was tagged with A1 tag for biotin or dye labeling. A structural model of mtIF2•GTPγS, fMet-tRNA$^{Met}$, and a leaderless mRNA bound on the mammalian mitochondrial ribosome (PDB 6GAW, with the 39S subunit hidden) placed this labeling site distal from the active sites of the ribosome. **(C)** Western blot results indicate homozygous tagging of the endogenous mS26 with the A1 tag. **(D)** Purified mS26-A1 tagged 28S subunits can be site-specifically labeled. **(E)** SDS-PAGE analysis of purified mtIF2, mtIF3, and bovine mtEF-Tu proteins used in this work. **(F)** Acid-urea PAGE analysis demonstrated efficiency charging of the human mitochondrial tRNA$^{Met}$. **(G)** The mS31 and bL31m labeling sites are highlighted on a structural model of the human mitochondrial 55S, indicating their distal locations from active sites of the ribosome. Western blots showing homogeneous tagging of mS31 (**H**) and bL31m (**I**) with the S6 tag. **(J)** The purified mS31-S6 tagged 28S subunit was site-specifically labeled with Cy3 (green) and the bL31m-S6 tagged 39S subunit labeled with Cy5 (red).

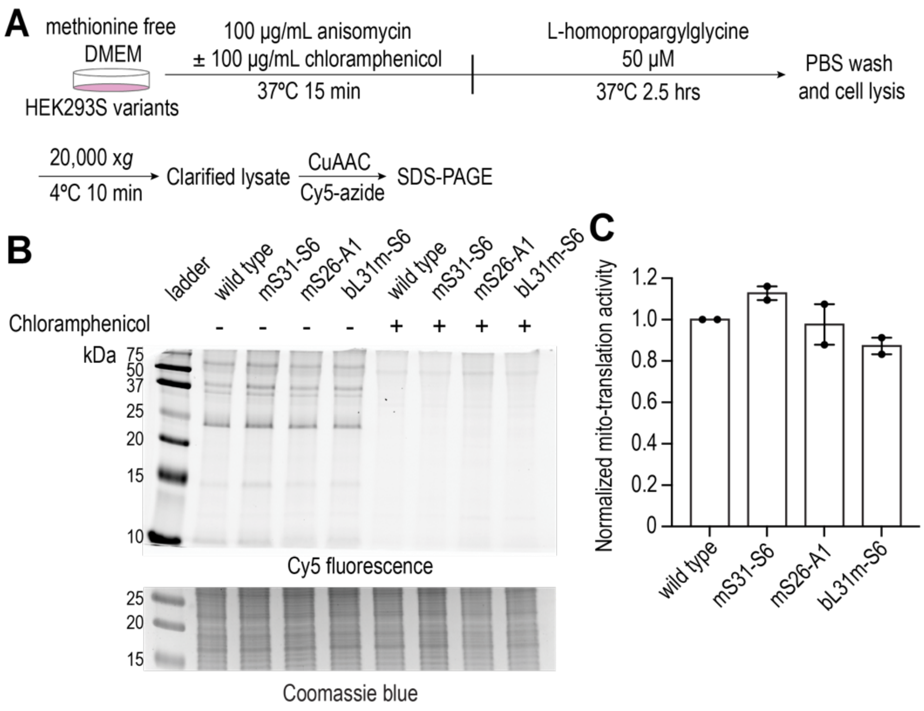

**Figure S3**. **Tagging of the mitoribosomes did not perturb their functions in vivo. (A)** Schematic of the on-gel Mito-FUNCAT assay. **(B)** SDS-PAGE results showing mitochondrial translation products (Cy5) with wild type and edited cells, which were suppressed when the cell was pre-treated with chloramphenicol. **(C)** Quantification of the mitochondrial translation activities. Indicated errors are standard error of the mean from two biological replicates.

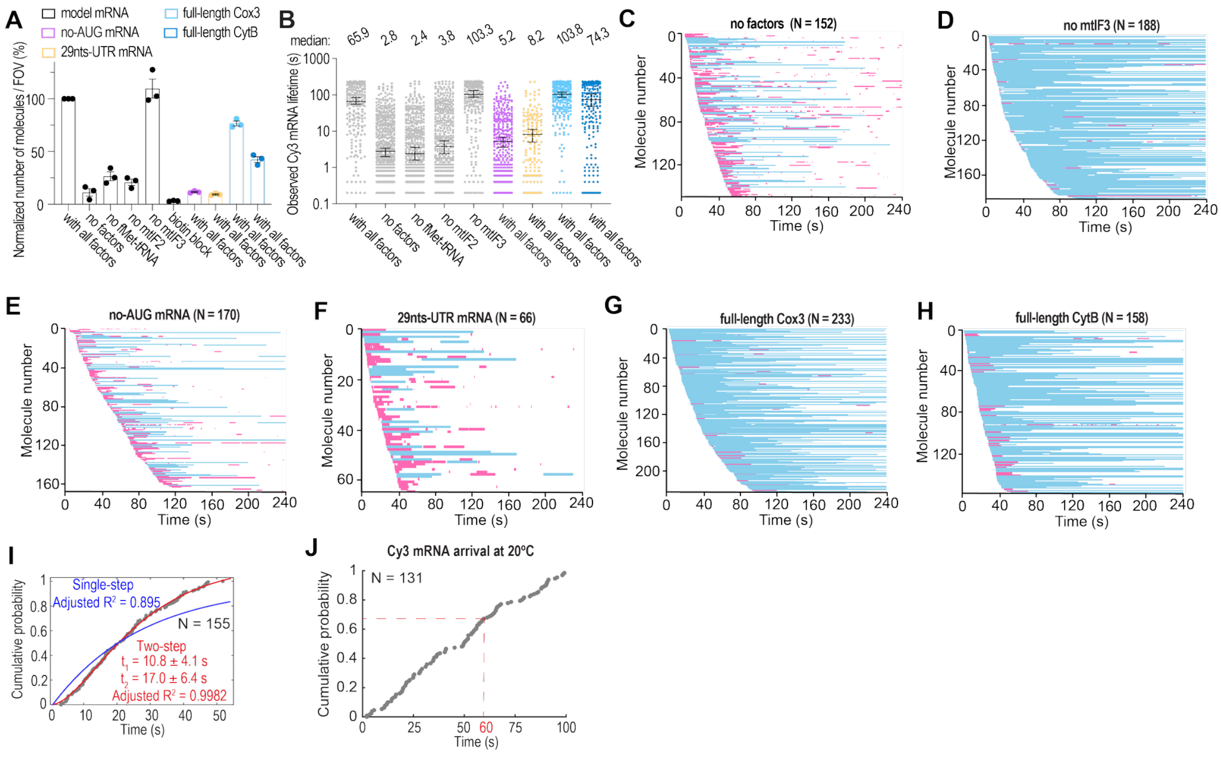

**Figure S4. Dynamics of mRNA binding to the 28S subunits. (A)** Mean normalized number of Cy3 molecules observed per field of view (FOV) in reactions under varying conditions. Indicated errors are standard error of the mean from three biological replicates with points representing the value from individual replicates. A part of the results is shown in Figure 1D. **(B)** Observed Cy3-mRNA binding lifetimes. Each point represents a single mRNA binding event. Red lines correspond to the median lifetimes (numbers shown on top), and errors are 95% confidence intervals of the median. Color codes are the same as in (**A**). **(C-H)** Stack of single-molecule traces from different reaction conditions, sorted by the arrival of the first Cy3 binding event. Each row represents a single 28S complex, and N represents the number of analyzed molecules. Cy3-mRNA binding events are displayed in blue (≥ 30 s) or pink (< 30 s). **(I)** Kinetics of Cy3-mRNA arrival to bind to the 28S subunit in the presence of all factors (gray dots). The red line shows the fit to a two-step kinetic model with the fitting results shown in red. The blue line shows the fit to a single-step reaction model with the fitting statistics results shown in blue. N represents the number of analyzed molecules. **(J)** Kinetics of Cy3 model mRNA arrival to bind to the 28S subunit in the presence of all factors at 20ºC (gray dots). N represents the number of analyzed molecules. The red dashed lines show the corresponding timepoint for the 28S PIC cryo-EM sample.

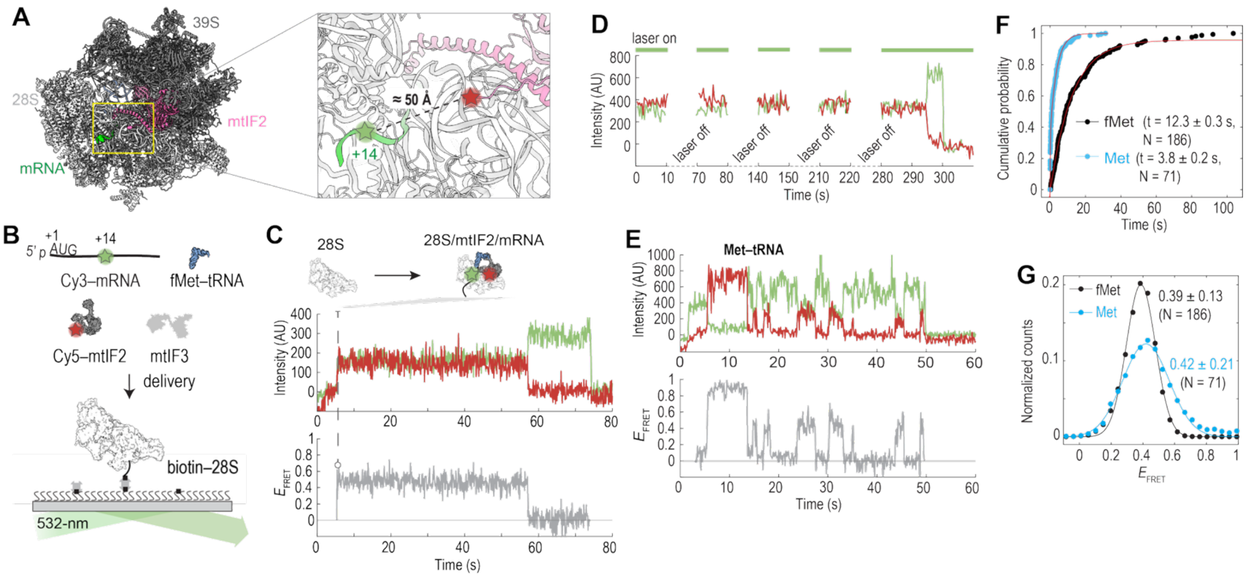

**Figure S5**. **Monitoring mRNA and mtIF2 binding to the 28S by FRET. (A)** The labeling sites on the mRNA and mtIF2 were within FRET distance. The 55S IC–mtIF2•GTPγS model (PDB 6GAW) is shown. **(B)** Schematic of assay to track Cy3-mRNA and Cy5-mtIF2 binding to tethered 28S subunits using a single 532-nm laser excitation. **(C)** Example fluorescence trace showing mtIF2 (Cy5, red) and mRNA (Cy3, green) binding to the 28S subunit. The Cy3 and Cy5 signals appeared simultaneously in a FRET state. However, any Cy5-mtIF2 events prior to the Cy3-mRNA binding would be missed in this experiment setup. **(D)** Time-lapse experiments showed that the lifetime of FRET between mtIF2 and mRNA was likely limited by dye photobleaching, suggesting long-lived binding of the mtIF2 and the mRNA on the 28S subunit. The surface was illuminated by the 532-nm laser for 10 s with 1 min intervals. **(E)** Example fluorescence trace showing mRNA (Cy3, green) binding to the 28S subunit followed by sampling of mtIF2 (Cy5, red) when Met-tRNA$^{Met}$ was used. **(F)** FRET lifetime distributions from experiments with fMet-tRNA$^{Met}$ (black) or Met-tRNA$^{Met}$ (blue) were fitted to a single-exponential model. The mean lifetimes (± 95% confidence interval) are shown, and N represents the number of analyzed molecules. As shown in (**D**), the FRET lifetime with fMet-tRNA$^{Met}$ was limited by the Cy5 dye photostability. **(G)** The mtIF2–mRNA FRET distributions in experiments with fMet-tRNA$^{Met}$ (black) or Met-tRNA$^{Met}$ (blue) were fit with a single-Gaussian distribution. Mean FRET efficiencies (± standard error of the mean) are shown, and N represents the number of analyzed molecules. The broader FRET distribution with Met-tRNA$^{Met}$ suggested dynamic heterogeneities in the complex.

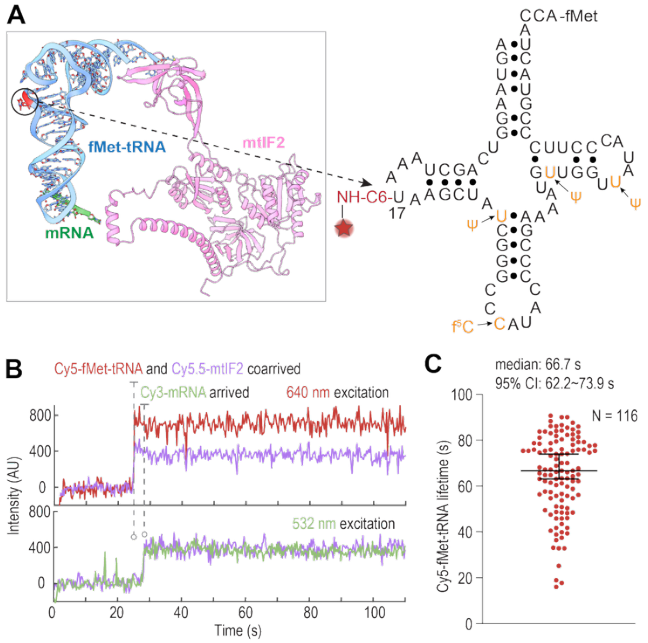

**Figure S6. Direct tracking of fMet-tRNA$^{Met}$. (A)** The U17 labeling site on the unmodified tRNA$^{Met}$ (yellow indicates the missing modifications) is flexible and distal from key interactions. **(B)** Example where mtIF2 (Cy5.5, purple) and fMet-tRNA$^{Met}$ (Cy5, red) co-arrived to bind the 28S subunit, followed by the mRNA (Cy3, green) binding, resulting in the mRNA–mtIF2 FRET. **(C)** Observed Cy5-fMet-tRNA$^{Met}$ binding lifetimes. Each point represents a single tRNA binding event. The median lifetime is shown with the 95% confidence interval. N represents the number of analyzed molecules.

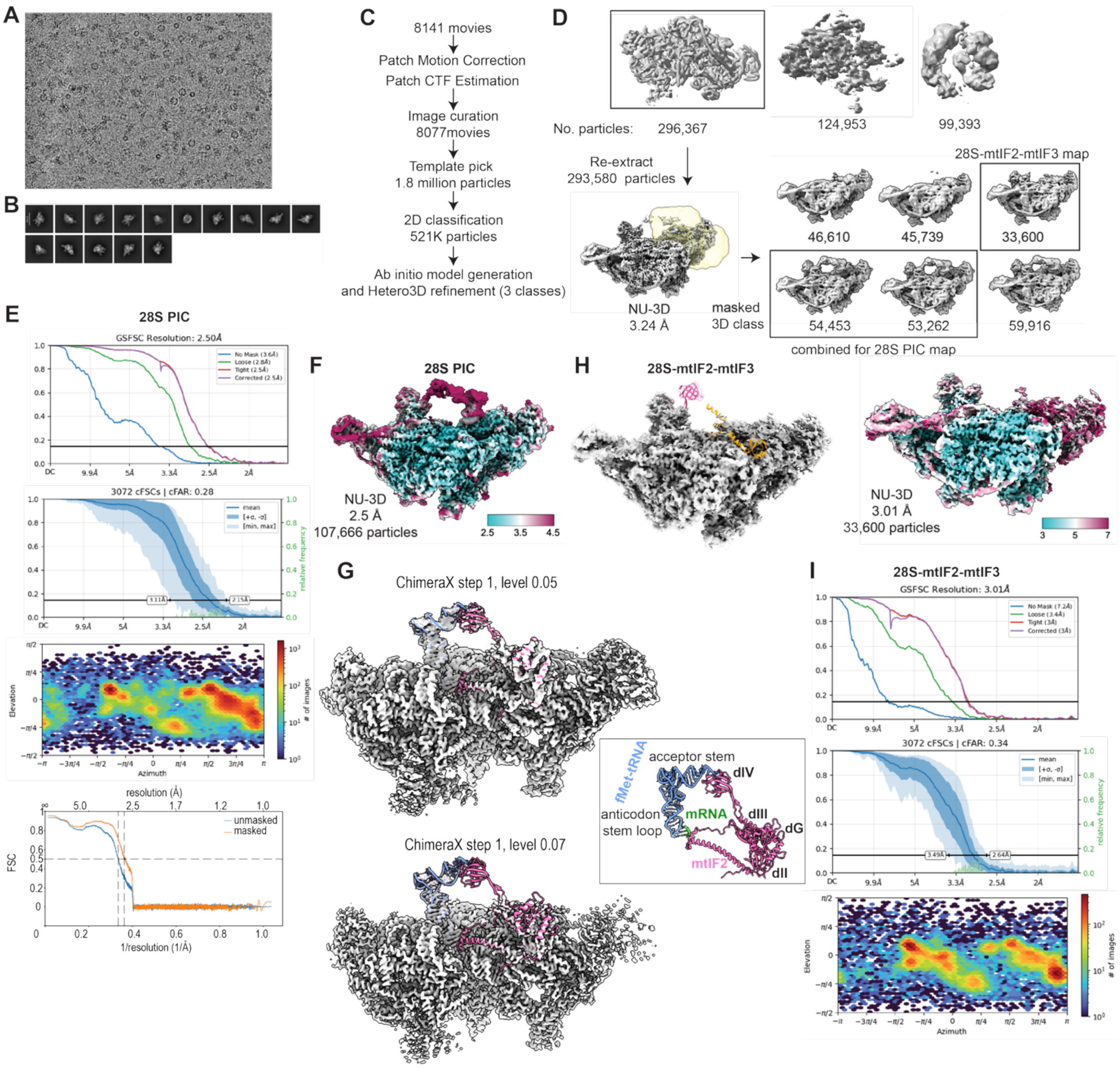

**Figure S7. Cryo-EM structure of the 28S PIC. (A)** Representative cryo-EM micrograph. **(B)** Representative two-dimensional (2D) class averages used in three-dimensional (3D) reconstruction. **(C)** Flow-chart for data processing. **(D)** Flow-chart for 3D classification and reconstruction. A 28S–mtIF2–mtIF3 complex and the 28S PIC were identified in the samples. **(E)** FSC curves, Angular distribution plot, and model-map FSC curves for the 28S PIC. **(F)** Local resolution distribution of the density map for the 28S PIC. **(G)** The 28S PIC map displayed at different contour levels, showing the heterogeneities in the peripheral regions of the fMet-tRNA$^{Met}$ and mtIF2. **(H)** The density map (left) and local resolution distribution (right) of the 28S–mtIF2–mtIF3 complex. PDB model 6RW5 was docked into the map, with mtIF2 shown in pink and mtIF3 in orange. **(I)** FSC curves and Angular distribution plot for the 28S–mtIF2–mtIF3 complex.

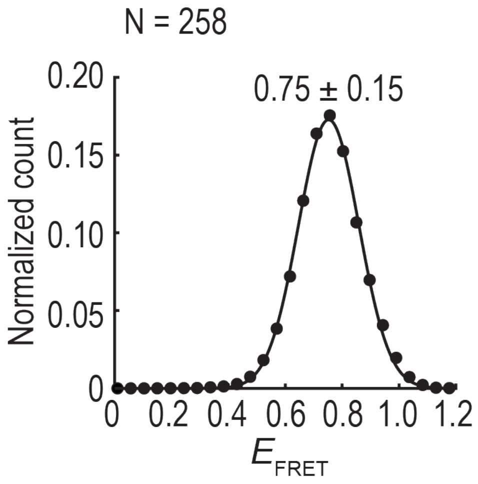

**Figure S8. Inter-ribosomal subunit FRET.** The 28S–39S FRET distribution in initiation experiments with the model mRNA was fit with a single-Gaussian distribution. Mean FRET efficiency (± standard error of the mean) is shown, and N represents the number of analyzed molecules. The observed high FRET efficiency was consistent with the predicted short distance between the two labeling sites on the subunits (~50 Å, Figure S2G).

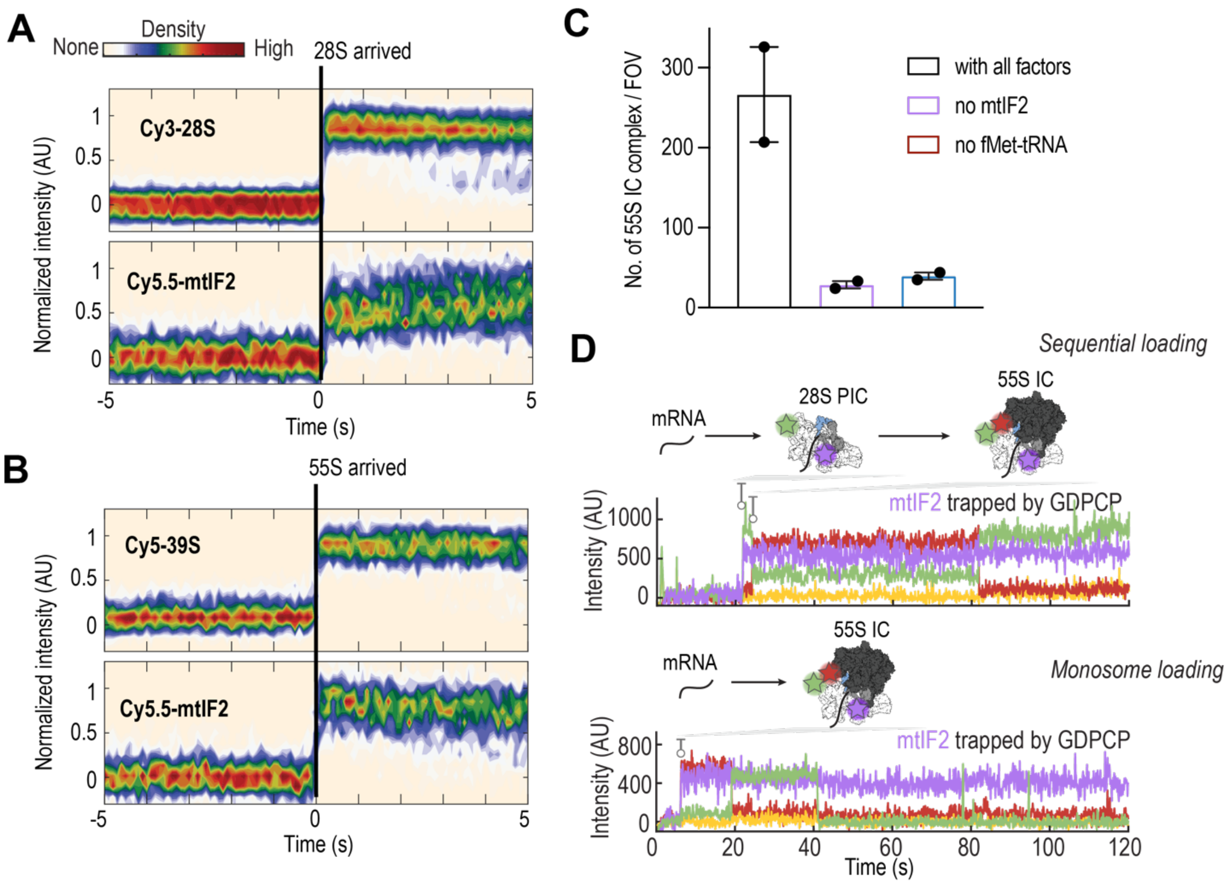

**Figure S9**. **Tracking mtIF2 throughout initiation and the transition to elongation. (A)** Density heat maps of the normalized fluorescence intensities of Cy5.5-mtIF2 and Cy3-28S synchronized to the binding of the 28S to the mRNA (N = 121). **(B)** Density heat maps of the normalized fluorescence intensities of Cy5.5-mtIF2 and Cy5-39S synchronized to the binding of the 39S to the mRNA (N = 65). The results in (**A**) and (**B**) indicated the co-arrival of mtIF2 with the 28S or 55S to bind the mRNA. **(C)** Both mtIF2 and fMet-tRNA$^{Met}$ were critical for initiation on leaderless mRNAs. The number of observed 55S IC assembled per FOV under different reaction conditions are shown. Indicated errors are standard error of the mean from two biological replicates with points representing the value from individual replicates. **(D)** Examples showing mtIF2 (purple) trapped on the 55S IC (28S in green and 39S in red) using GDPCP in both the sequential loading (upper) and monosome loading (bottom) initiation pathways, preventing Phe-TC (yellow) from binding to the ribosomal A site.

**Table S1. DNA oligos used in this study.**

| Oligo ID | Description | Sequence |
|---|---|---|
| JW92 | mS31 guide-forward | CACCGATACAGTTCAATTAAGACCA |
| JW93 | mS31 guide-reverse | AAACTGGTCTTAATTGAACTGTATC |
| JW94 | mS31-S6 repair template | GTTTAGAAATTATTTTAATGAAAAAAAGGATATTCTAAAAGAAAGTAACATACAGTTCAATGGCGATTCTCTTAGTTGGCTGTTGCGTCTTCTGAACTAAGACCATCGAAATTTTTATTTCAAACAATTAGAGATGGATATTACAACTAAATAAAATAATTTTACTAG |
| JW95 | Screening PCR primer for mS31 editing | GTAATGTATTTGGTCTCATTTCCTAG |
| JW96 | Screening PCR primer for mS31 editing | CAATGTAACATTTATACCAGCC |
| JW100 | bL31m guide-forward | CACCGTCCCGGGGTGGCTGGAGCCA |
| JW101 | bL31m guide-reverse | AAACTGGCTCCAGCCACCCCGGGAC |
| JW102 | bL31m-S6 repair template | GAACGAGTGATTTAAGAACAGCCCTCCCATCTTAGCAATGTCCCGGGGTGGCTGGAGCCACCGTCAGTTCAGAAGACGCAACAGCCAACTAAGAGAATCGCCCTTCTTGGTCCTGGTCCAGAACTGTCGGTAGCGCTCCACATGCAAGTCATCACTGAGCTCCTGCTCGTAC |
| JW103 | Screening PCR primer for bL31m editing | CCAGGTGGCAGCTTCAAG |
| JW104 | Screening PCR primer for bL31m editing | GAAGGGAGAGACCAATAAGAATG |
| JW747 | mS26 guide-forward | CACCG GGACTCCTAGGGGCCCAGTA |
| JW748 | mS26 guide-reverse | AAAC TACTGGGCCCCTAGGAGTCC C |
| JW749 | mS26-A1 repair template | GGGCCATCACCAGAGAGGGGCTGGTGGTCAGGCCACAACGCAGGGACTCCGGCGATTCTCTTGATATGTTGGAATGGTCTCTGATGTAGGGGCCCAGTAAGGACAGTGCCCGCCAGGGACCATGTATGTATCATGG |
| JW750 | Screening PCR primer for mS26 editing | CTTCATCACCCGAGAGAAC |
| JW751 | Screening PCR primer for mS26 editing | CTGCTTTATTCCAGGTCAGG |
| JW445 | PCR amplification of full-length Cox3 or CytB mRNAs_forward | GTGGAATTGTGAGCGGATAAC |
| JW446 | PCR amplification of full-length CytB mRNA_reverse | TTTTTTTTTTTTTTTTTTTTTTTTTTTTTTTTTTTTTTAGGCCCATTTGAGTATTTTG |
| JW447 | PCR amplification of full-length Cox3 mRNA_reverse | TTTTTTTTTTTTTTTTTTTTTTTTTTTTTTTTTTTTTTAAGACCCTCATCAATAG |
| JW131 | Primer for PCR to introduce the ybbR tag sequence to the mtIF2 expression plasmid | TCGCCTCAAAGTTGGCATGGAAAAGCCAGCTGAGCAG |
| JW141 | Primer for PCR to introduce the ybbR tag sequence to the mtIF2 expression plasmid | TGCCAACTTTGAGGCGATGAACTCCAAGCTATCGCCGCTACCGCTGCCAC |
| JW13 | TIRF experiment blocking oligo 1 | CGTTTACACGTGGGGTCCCAAGCACGCGGCTACTAGATCACGGCTCAGCT |
| JW14 | TIRF experiment blocking oligo 2 | AGCTGAGCCGTGATCTAGTAGCCGCGTGCTTGGGACCCCACGTGTAAACG |

### Table S2. Cryo-EM data collection, refinement and validation statistics

| | 28S PIC | 28S-mtIF2-mtIF3 |
|---|---|---|
| **Data collection and processing** | | |
| Magnification | 105,000 | 105,000 |
| Voltage (kV) | 300 | 300 |
| Electron exposure (e–/Å$^2$) | 50 | 50 |
| Defocus range (μm) | -1.2 to -2.5 | -1.2 to -2.5 |
| Pixel size (Å) | 0.826 | 0.826 |
| Symmetry imposed | - | - |
| Initial particle images (no.) | 296,367 | 296,367 |
| Final particle images (no.) | 107,666 | 33,600 |
| Map resolution (Å) | 2.5 | 3.01 |
| FSC threshold | 0.143 | 0.143 |
| Map resolution range (Å) | 2.5-5 | 3-7 |
| **Refinement** | | |
| Initial model used (PDB code) | 7PO2 | N/A |
| Model resolution (Å) | 2.8 | |
| FSC threshold | 0.5 | |
| Model resolution range (Å) | - | |
| Map sharpening *B* factor (Å$^2$) | - | |
| Model composition | | |
| Non-hydrogen atoms | 76047 | |
| Protein residues | 6304 | |
| Nucleotide | 1031 | |
| Ligands | GTP:1 | |
| | ZN: 1 | |
| | FES: 1 | |
| | ATP: 1 | |
| | K: 20 | |
| | MG: 72 | |
| *B* factors (Å$^2$) | | |
| Protein | 105.88 | |
| Nucleotide | 80.59 | |
| Ligand | 86.26 | |
| R.m.s. deviations | | |
| Bond lengths (Å) | 0.005 | |
| Bond angles (°) | 0.566 | |
| Validation | | |
| MolProbity score | 1.30 | |
| Clashscore | 5.28 | |
| Poor rotamers (%) | 0.00 | |
| Ramachandran plot | | |
| Favored (%) | 97.95 | |
| Allowed (%) | 2.02 | |
| Disallowed (%) | 0.03 | |
